# Heme oxygenase-2 (HO-2) binds and buffers labile heme, which is largely oxidized, in human embryonic kidney cells

**DOI:** 10.1101/2021.06.06.447256

**Authors:** David A. Hanna, Courtney M. Moore, Liu Liu, Xiaojing Yuan, Angela S. Fleischhacker, Iqbal Hamza, Stephen W. Ragsdale, Amit R. Reddi

## Abstract

Heme oxygenases (HO) detoxify heme by oxidatively degrading it into carbon monoxide, iron, and biliverdin, which is reduced to bilirubin and excreted. Humans express two isoforms: inducible HO-1, which is up-regulated in response to various stressors, including excess heme, and constitutive HO-2. While much is known about the regulation and physiological function of HO-1, comparatively little is known about the role of HO-2 in regulating heme homeostasis. The biochemical necessity for expressing constitutive HO-2 is largely dependent on whether heme is sufficiently abundant and accessible as a substrate under conditions in which HO-1 is not induced. By measuring labile heme, total heme, and bilirubin in human embryonic kidney HEK293 cells with silenced or over-expressed HO-2, and various HO-2 mutant alleles, we found that endogenous heme is too limiting to support HO-2 catalyzed heme degradation. Rather, we discovered that a novel role for HO-2 is to bind and buffer labile heme. Taken together, in the absence of excess heme, we propose that HO-2 regulates heme homeostasis by acting as a heme buffering factor in control of heme bioavailability. When heme is in excess, HO-1 is induced and both HO-2 and HO-1 can provide protection from heme toxicity by enzymatically degrading it. Our results explain why catalytically inactive mutants of HO-2 are cytoprotective against oxidative stress. Moreover, the change in bioavailable heme due to HO-2 overexpression, which selectively binds ferric over ferrous heme, is consistent with the labile heme pool being oxidized, thereby providing new insights into heme trafficking and signaling.

## Introduction

Heme is an essential but potentially cytotoxic metallocofactor and signaling molecule (1–10). Consequently, cells must tightly regulate the concentration and bioavailability of heme (8,11–14). In mammals, the total intracellular concentration of heme is governed by the relative rates of *de novo* synthesis, degradation, import, and export. The atomic resolution structures and chemical mechanisms of all the heme biosynthetic and catabolic enzymes are known and well understood (11, 15). While cell surface heme importers (16) and exporters (17, 18) have been identified, their molecular mechanisms remain poorly characterized and, outside of developing red blood cells in the case of heme exporters, the physiological context in which they function is unclear and controversial (19). The bioavailability of heme, which is comparatively less well understood, is governed by a poorly characterized network of heme buffering factors, intracellular transporters, and chaperones that ensure heme is made available for heme dependent processes located throughout the cell.

When cells are confronted with excess heme, heme synthesis is down-regulated (11,20,21) and heme can be detoxified by storage into lysosome-related organelles (22, 23), export (24, 25), or degradation (26–31). Arguably, the best understood mechanism for heme detoxification is through the heme catabolism pathway. The first and rate-limiting step of heme degradation is catalyzed by the heme oxygenases (HO) (32, 33). Mammals encode two HO isoforms, inducible HO-1 and constitutive HO-2 (34–37). HO-1 and HO-2 are structurally similar, both in primary sequence and tertiary structure, operate using the same chemical mechanism, and exhibit similar catalytic properties, including Michaelis constants (*K*_M_) and maximal velocities (*V*_max_) (37–39). HOs, which are primarily anchored into the endoplasmic reticulum (ER) membrane and whose active sites face the cytoplasm, bind oxidized ferric heme in its resting state using a histidine axial ligand. Upon reduction, using electrons from the NADPH-cytochrome P450 reductase (CPR) system, and dioxygen binding (O_2_), HOs catalyze the oxidative degradation of heme to form biliverdin, ferrous iron (Fe^2+^), and carbon monoxide (CO) (40–44). Biliverdin is subsequently rapidly metabolized to bilirubin via a NADPH-biliverdin reductase and expelled from cells (45, 46). Given that heme catabolites Fe^2+^, CO, biliverdin, and bilirubin have their own distinct beneficial or detrimental effects on cell physiology in various contexts, the activity of HO enzymes and availability of its heme substrate can impact metabolism in numerous ways (32,47–55).

While the structures and mechanisms of HO-1 and HO-2 are largely the same (37-39,41,43,56), the regulation and expression of these two enzymes is very different (27,31,35). HO-1, which is comparatively far better understood, is induced by excess heme, as well as several non-heme stressors like oxidative stress, infection, and exposure to various xenobiotics (57–61). HO-2, on the other hand, is constitutively expressed across all tissues and cell types, being most abundant in the brain and testis (34, 35). The current rationale for dual mammalian HO isoforms is that HO-2 provides a baseline level of protection from heme in the absence of cellular stressors that would otherwise induce HO-1. However, the biochemical necessity for expressing constitutive HO-2 is largely dependent on whether sufficient heme is available as a substrate under conditions in which HO-1 is not induced.

Total cellular heme in yeast and various non-erythroid human cell lines is on the order of 1-20 µM (62–67). All heme in the cell partitions between exchange inert high affinity hemoproteins, such as cytochromes and other heme enzymes, and certain exchange labile heme complexes that buffer free heme down to nanomolar concentrations (8,12–14,63,68,69). The factors that buffer heme are poorly understood, but likely consist of a network of heme binding proteins, nucleic acids, and lipid membranes (8,12–14,63,66,67,70,71). Labile heme (LH) may act as a reservoir for bioavailable heme that can readily exchange with and populate heme binding sites in heme dependent or regulated enzymes and proteins. The nature of LH, including its speciation, oxidation state, concentration and distribution are not well understood, but may be relevant for the mobilization and trafficking of heme. It is currently not known what the source of heme is for HOs, *i.e.* whether it is buffered free heme, certain labile heme complexes, or a dedicated chaperone system that traffics and channels heme to HO in a manner that bypasses the LH pool.

The recent development of fluorescence and activity-based heme sensors has offered unprecedented insights into labile heme and their diverse roles in physiology (63,68,69,72,73). Strictly speaking, these probes report on the *availability* of heme to the sensor, not necessarily *free* heme coordinated by water (8,13,74). In other words, the heme occupancy of the sensor is dictated by the extent to which the labile heme pool can exchange with the probe. However, many investigators convert the fractional heme loading of a probe to a buffered free heme concentration, which can be done if the heme-sensor dissociation constant is known. While problematic in that the sensor may not be probing “free heme”, it nonetheless provides a measure of labile heme since the calculated concentration of free heme is related to sensor heme occupancy.

In intact living yeast and various non-erythroid human cell lines, estimates of buffered free heme based on genetically encoded heme sensors are on the order of ∼5-20 nM (63,68,69). If free heme is a heme source for HO-2, which has a *K*_M_ value of 400 nM for heme (37), it is expected to be less than 5% active (**Figure 1**, black curve), assuming other HO substrates, *e.g.* CPR and O_2_ are not limiting. In contrast, using heme reporters in human embryonic kidney HEK293 and human lung fibroblast IMR90 cell extracts, it was found that free heme was as high as 400-600 nM (73, 75), corresponding to heme concentrations in which HO-2 is ∼50% active (**Figure 1**, black curve). Given the relatively large span in estimates of free heme using different detection methods, between 20 and 600 nM, it is not clear how active HO-2 is, raising the intriguing possibility that it may have alternative roles in heme cell biology distinct from enzymatic heme degradation. Indeed, prior reports have found that enzymatically inactive HO-2 alleles can rescue cells from oxidative stress (76). Alternatively, if heme were delivered to HO-2 using a specific heme delivery system that bypasses LH, the activity of HO-2 would be dependent on access to such heme chaperones, making LH or buffered free heme irrelevant towards understanding the activity of HO-2 in cells.

**Figure 1.**
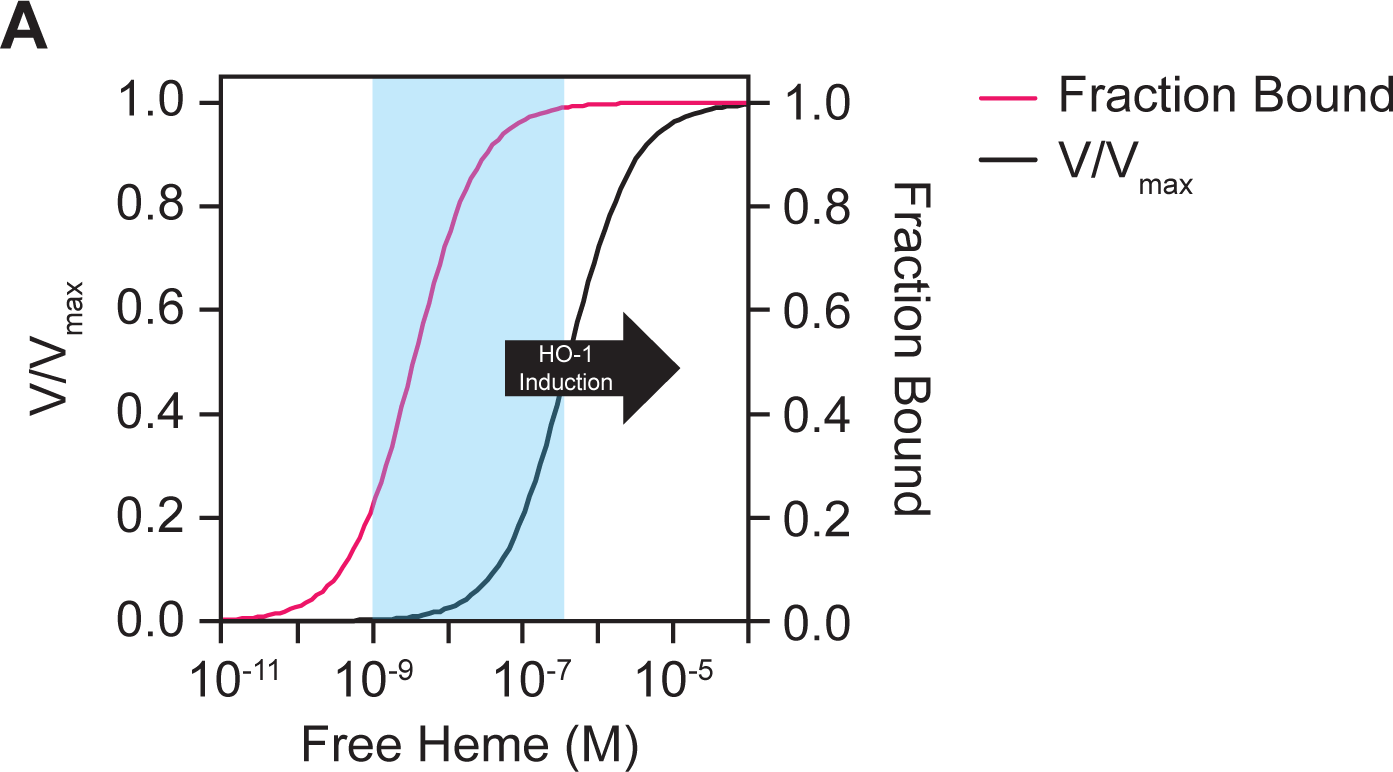
Fractional heme occupancy and activity of HO-2 as a function of free heme concentration. The heme occupancy of HO-2 (red curve; right y-axis) was simulated using a 1- site heme binding model governed by the following equation: [Heme] / {[Heme] + *K*_D_}, assuming a heme-HO-2 *K*_D_ value of 3.6 nM. The fraction of maximal velocity, V/V_max_, (black curve; left y-axis) was simulated using a Michaelis-Menten model governed by the following equation: [Heme] / {[Heme] + *K*_M_}, assuming a heme *K*_M_ value of 400 nM. The light blue shaded region represents the span of values reported for buffered free heme in cells and the black arrow indicates the concentration of intracellular free heme in which HO-1 is induced.

An alternative model for constitutive HO-2 function is that it acts as a component of a larger network of proteins that buffer heme. The ferric heme dissociation constant (*K*_DIII_) for HO-2 is 3.6 nM (77) and, at the lower estimates of buffered free heme concentrations, is expected to be fractionally populated with heme (**Figure 1**, red curve). As such, HO-2 may act in a larger network of exchange labile heme complexes and serve as an access point for heme distribution, possibly to the ER. Such a model would require that LH is largely oxidized and exchangeable with HO-2 and would predict that perturbations in HO-2 expression will perturb LH or buffered free heme.

In the present report, using the model human cell line, HEK293 cells, we sought to determine if endogenous LH is sufficiently accessible and abundant for HO-2 to be actively catabolizing heme and establish the oxidation state of LH. HEK293 cells were chosen because of extensive prior work in these cell lines probing HO-2 function (64,76,78) and labile heme (67-69,73). By measuring labile heme, total heme, and bilirubin in HEK293 cells with silenced or over-expressed HO-2, and various mutant HO-2 alleles, we found that heme is accessible to HO- 2, but too limiting to support a role for HO-2 in actively degrading heme. Rather, our data support a role for HO-2 in regulating heme bioavailability by buffering it. Moreover, the change in LH due to HO-2 overexpression is consistent with LH being largely oxidized. Altogether, our findings force us to re-think the physiological role of constitutive HO-2 in cell types that have limiting levels of LH to support its catalytic activity.

## Results

### Measuring labile heme in HEK293 cells

In order to determine how active HO-2 is in HEK293 cells, we first sought to determine the amount of cytosolic labile heme. Towards this end, we transiently transfected previously described genetically encoded fluorescent heme sensors into HEK293 cells and analyzed labile heme levels using flow cytometry. Heme sensor 1 (HS1) is a tri-domain construct consisting of a heme binding domain, cytochrome *b*_562_ (Cyt *b*_562_), an enhanced green fluorescent protein (eGFP) whose emission is quenched by heme, and a red fluorescent protein (mKATE2) whose emission is relatively unaffected by heme (63). Thus, the eGFP/mKATE2 fluorescence ratio of HS1 decreases upon heme binding and is a reporter of the buffered exchange labile or bioavailable heme pool in cells.

We analyzed intracellular labile heme (LH) using the high affinity heme sensor, HS1, which binds heme using Cyt *b*_562_ Met_7_ and His_102_, a moderate affinity heme sensor, HS1-M7A, and a variant that cannot bind heme, HS1-M7A, H102A (63). Between pH 6.8 – 7.4, the consensus range of cytosolic pH values reported for HEK293 cells (79–81), the ferric and ferrous heme dissociation constants for HS1 are *K*_DIII_ = 3 nM and *K*_*D*_^II^ = 1 pM and for HS1-M7A are *K*_DIII_ = 1 μM and *K*_*D*_^II^ = 25 nM (62, 63). As shown in **Figures 2a** and **2b**, flow cytometric analysis of HS1 eGFP/mKATE2 fluorescence ratios indicate that it is heme responsive. Compared to cells cultured in regular media (Reg Media), HS1 expressed in HEK293 cells depleted of intracellular heme by culturing in heme deficient (HD) media and with the heme biosynthetic inhibitor succinylacetone (SA) for 24 hours have a characteristically high median eGFP/mKATE2 fluorescence ratio. In contrast, cells cultured with 10 μM heme for 24 hours have a characteristically low median eGFP/mKATE2 fluorescence ratio (**Figures 2a** and **2b**). On the other hand, HS1-M7A and HS1-M7A, H102A are not heme responsive since eGFP/mKATE2 fluorescence ratios were not sensitive to heme depletion (HD+SA) or heme excess (10 μM heme) (**Figures 2a** and **2b**). *In toto*, assessment of the median eGFP/mKATE2 fluorescence ratios from replicate flow cytometry data (**Figure 2b**) indicate that HS1 binds heme tightly enough to measure endogenous LH in cells, but HS1-M7A does not.

**Figure 2.**
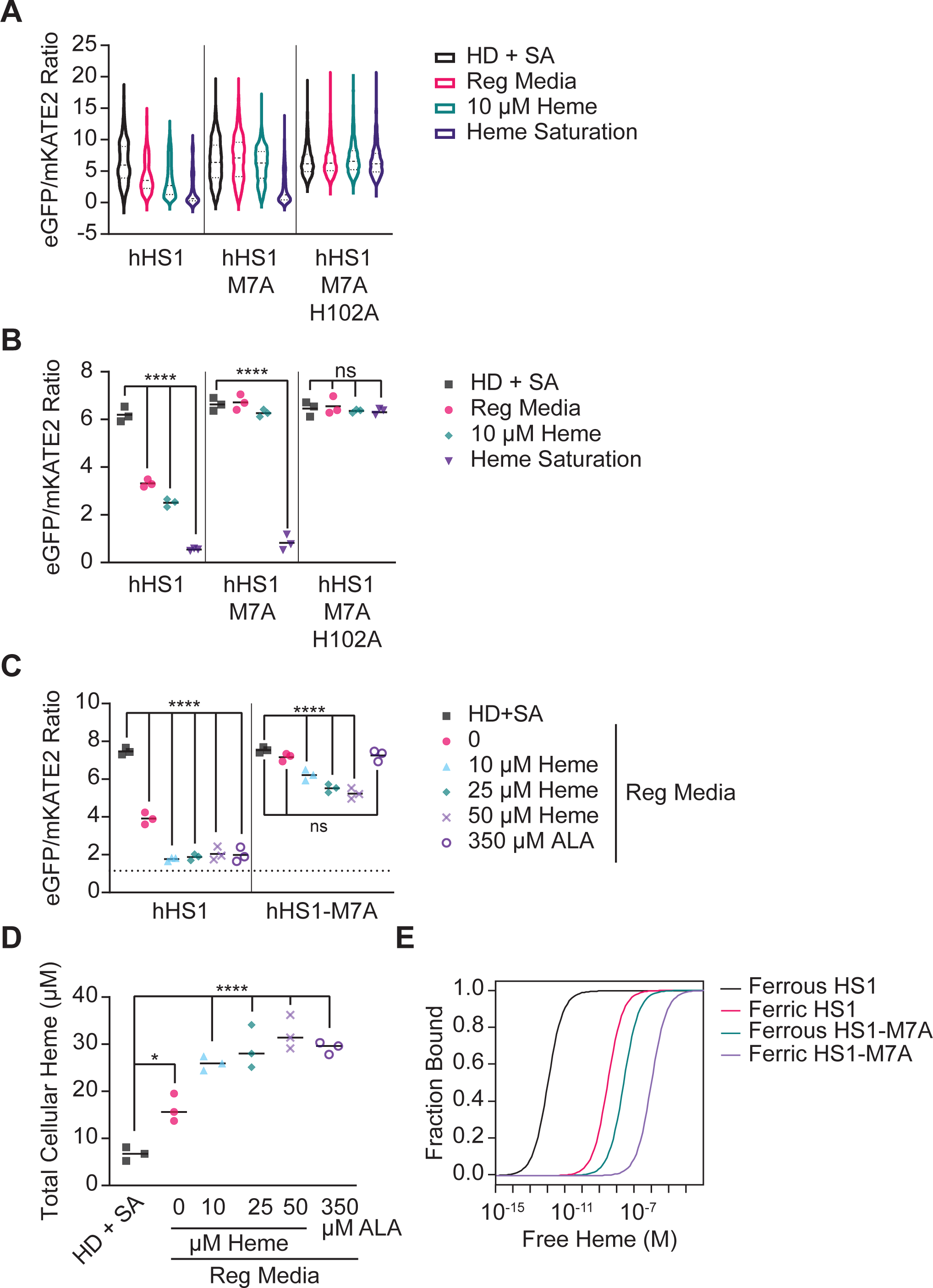
Heme Sensor 1 (HS1) can sense labile heme in HEK293 cells. (**a**) Representative violin plots depicting the distribution of eGFP/mKATE2 fluorescence ratios in single HEK293 cells expressing the high affinity heme sensor, HS1, the moderate affinity heme sensor, HS1-M7A, and the heme binding incompetent control scaffold, HS1-M7A, H102A. The heme responsiveness of the sensors was established by culturing HEK293 cells in regular media (Reg. Med.; DMEM with 10% FBS), heme deficient (HD) media supplemented with succinylacetone (SA) (HD + SA; DMEM with heme depleted 10% FBS and 500 µM SA), or in regular media with 10 µM hemin chloride (10 µM Heme). In order to saturate the sensors, cells were permeabilized with digitonin and incubated with 100 µM hemin chloride (Heme Saturation; serum free DMEM with 1 mM ascorbate, 40 µM digitonin, 100 µM hemin chloride). (**b**) Median heme sensor eGFP/mKATE2 fluorescence ratio values derived from flow cytometry experiments (as in panel **a**) from triplicate HEK293 cultures. (**c**) Median heme sensor eGFP/mKATE2 fluorescence ratio values derived from flow cytometry experiments from triplicate HEK293 cultures grown in HD + SA media or in regular media supplemented with the indicated concentrations of hemin chloride or 5-aminolevulinic acid (ALA) for 24 hours. Representative violin plots of eGFP/mKATE2 fluorescence ratio distributions from single cell analysis of HEK293 cultures are shown in **Figure S8a**. (**d**) Measurements of total heme in triplicate HEK293 cultures grown in HD + SA media or in regular media supplemented with the indicated concentrations of hemin chloride or 5-aminolevulinic acid (ALA) for 24 hours. (**e**) Relationship between sensor heme occupancy for HS1 and HS1-M7A and buffered free heme, depending on weather heme is oxidized (ferric) or reduced (ferrous). See main text and **Experimental Procedures** for details. The statistical significance is indicated by asterisks using one-way ANOVA for multiple comparisons using Tukey’s range test. *p<0.05, **p<0.01,***p <0.001, ****p<0.0001, ns = not significant.

In order to better quantify LH, we adapted a previously developed sensor calibration protocol to relate the sensor fluorescence ratios to its heme occupancy (63). The fractional saturation of the heme sensor is governed by **Equation 1** (63):

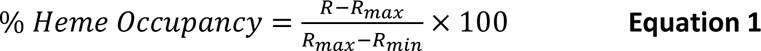

where R is the eGFP/mKATE2 fluorescence ratio under any given experimental condition, R_max_ is the eGFP/mKATE2 fluorescence ratio when the sensor is saturated with heme, and R_min_ is the eGFP/mKATE2 fluorescence ratio when the sensor is depleted of heme. R_max_ is determined by permeabilizing cells with digitonin and adding an excess of heme to drive heme binding to the sensor (“Heme Saturation” in **Figures 2a** and **2b**). R_min_ is determined by growing cells with succinylacetone (SA) in media that was depleted of heme (HD) (“HD + SA” in **Figures 2a** and **2b**). Analysis of the median fluorescence ratios from replicate flow cytometry measurements (**Figure 2b**) indicate that the heme occupancies of HS1 and HS1-M7A are ∼50% and ∼0%, respectively, when HEK293 cells are cultured in regular media.

Cells that accumulate excess intracellular heme, up to 30 μM, by supplementation with 350 μM 5-aminolevulinic acid (ALA), a heme biosynthetic precursor, to induce heme synthesis or with 10 - 50 μM exogenous heme, saturate HS1 to nearly ∼100% heme occupancy (**Figures 2c** and **2d**). As a consequence, ALA supplementation can also be used as a calibration control to saturate HS1 in control experiments to establish % heme occupancy. In contrast, HS1-M7A heme occupancy remains at ∼0% with ALA supplementation and only increases to ∼30% fractional heme saturation when intracellular heme approaches 30 μM with exogenous heme supplementation (**Figures 2c** and **2d**).

If one assumes the sensor is binding *free* heme, the concentration of buffered free heme can be determined by **Equation 2** (63):

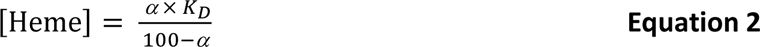

where *K*_D_ is the heme sensor-heme dissociation constant and α is the heme occupancy of the sensor defined by **Equation 1**. If one assumes LH is largely reduced, the buffered free heme concentration can be estimated to be ∼ 1 pM in HEK293 cells cultured in regular media by assuming the HS1 median R_min_, R_max_ and R values from **Figure 2b** or **2c**, as well the HS1 *K*_*D*_^II^ value of 1 pM. On the other hand, if one assumes LH is largely oxidized, the buffered free heme concentration can be estimated to be ∼5 nM, assuming a HS1 *K*_*D*_^II^ value of 3 nM and the aforementioned median R_min_, R_max_ and R values from **Figures 2b** or **2c**.

Uncertainties in the concentration of free heme stems from a lack of knowledge of the specific oxidation state of LH, which is addressed further below, and a means to precisely determine R_min_. The variation in the concentrations of free heme can be estimated based on the HS1 and HS1-M7A heme occupancies, their ferric or ferrous heme dissociation constant values, and assumptions about the oxidation state of LH (**Figure 2e**). Regarding R_min_, since heme is absolutely required for human cell lines, heme depletion by culturing cells in HD+SA media does not completely eliminate intracellular heme and results in only a 2-fold decrease compared to cells cultured in regular media (**Figure 2d**) (64). Past studies in *Saccharomyces cerevisiae*, which can be cultured in the complete absence of heme if supplemented with oleic acid and ergosterol, found that labile heme is depleted by > 90% when total heme is diminished by only 2-fold due to SA treatment (62). The degree to which HEK293 cells are yeast-like in the sensitivity of their LH pools to heme depletion will dictate the accuracy of R_min_ and therefore sensor heme occupancy and [LH]. Irrespective of the precise values of buffered free heme and R_min_, or if free heme even exists, the eGFP/mKATE2 fluorescence ratios of HS1, relative to *observed* R_min_ and R_max_ values, can nonetheless be used as a readout of sensor heme loading and exchange labile heme.

### HO-2 regulates heme bioavailability but not heme degradation in HEK293 cells

Having established that we can probe LH in HEK293 cells using HS1, we next sought to determine the effects of HO-2 silencing on the intracellular concentrations of labile heme, total heme, and bilirubin, a readout of HO activity. Silencing HO-2 using small interfering RNA (siRNA) resulted in ∼80% depletion of steady-state HO-2 levels and did not alter HO-1 expression (**Figure 3a**). HO-2 silencing results in a small but statistically significant increase in LH, with the median eGFP/mKATE2 fluorescence ratio decreasing from 1.1 to 0.9, which corresponds to a 11% increase in HS1 heme occupancy, from 63% to 74% (**Figure 3b**). In contrast, the cellular heme degrading activity of HOs are not affected by HO-2 as evidenced by total heme (**Figure 3c**) and bilirubin levels (**Figure 3d**) being unchanged by ablating HO-2. Heme degrading activity only increases when cells are challenged with elevated intracellular heme. Cells cultured with 350 μM ALA, which increases total and labile heme (**Figure 2d**), exhibit significantly elevated bilirubin levels (**Figure 3d**). Cells cultured with ALA have elevated HO-1 expression, up to ∼10- fold, with no change in HO-2 levels (**Figure 3a**). In total, our data indicate that HO-2 expression affects LH but not heme degradation in HEK293 cells.

**Figure 3.**
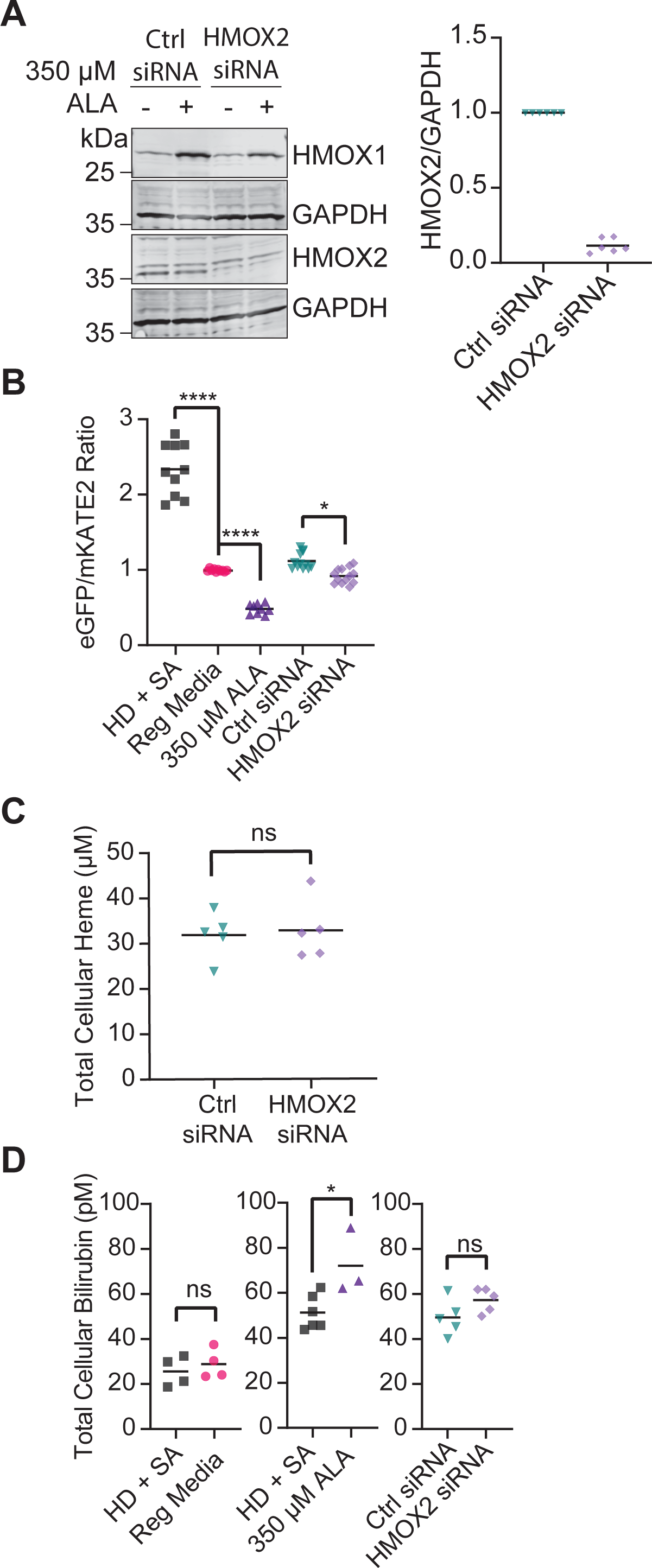
HO-2 silencing increases labile heme but does not affect total heme or bilirubin levels. (**a**) Representative immunoblot of HO-1 (HMOX1), HO-2 (HMOX2), and GAPDH expression in HEK293 cells treated with or without 350 µM 5-aminolevulinic acid (ALA) and scrambled (Ctrl siRNA) or targeted (HMOX2 siRNA) siRNA against HO-2 (Left). Immunoblot analysis from 6 independent trials demonstrates that siRNA against HMOX2 results in ∼80-90% silencing of HO- 2 protein expression (Right). (**b**) Median heme sensor eGFP/mKATE2 fluorescence ratio values derived from flow cytometry experiments from 9-13 replicates of HEK293 cultures grown in HD + SA media, regular media, or regular media supplemented with 350 µM ALA or control or targeted siRNA against HMOX2. Representative violin plots of eGFP/mKATE2 fluorescence ratio distributions from single cell analysis of HEK293 cultures are shown in **Figure S8b**. (**c**) Measurements of total heme in quintuplicate HEK293 cultures grown in regular media supplemented with control or targeted siRNA against HMOX2. (**d**) Measurements of total bilirubin in 3-6 replicates of HEK293 cultures grown in in HD + SA media, regular media, or regular media supplemented with 350 µM ALA or control or targeted siRNA against HMOX2. The statistical significance is indicated by asterisks using one-way ANOVA for multiple comparisons using Tukey’s range test. *p<0.05, **p<0.01,***p <0.001, ****p<0.0001, ns = not significant.

### HO-2 over-expression decreases heme availability in a manner that does not require its catalytic activity

Having established that HO-2 depletion increases LH, we sought to determine if HO-2 overexpression could decrease it. Towards this end, an HO-2 overexpression plasmid was transiently transfected into HEK293 cells, resulting in greater than 10-fold increase in HO-2 expression, without affecting HO-1 expression (**Figure 4a**). Moreover, ALA-induced overproduction of heme does not alter steady-state HO-2 levels like it does with HO-1, which is induced by heme (**Figure 4a**). HO-2 overexpression does not affect total heme in either heme- deplete or replete conditions (**Figure 4b**) or alter bilirubin levels (**Figure 4c**). To rule out the possibility that electron delivery to the heme oxygenase system via CPR may limit heme oxygenase activity (76, 78), we also assessed bilirubin production upon addition of 350 μM ALA to increase heme synthesis and found that HO-2 overexpressing cells had a concomitant increase in bilirubin levels (**Figure 4c**). Interestingly, HO-2 overexpression significantly depletes LH in the cytosol, resulting in a shift in HS1 heme occupancy from ∼55% to 0% (**Figure 4d**). Using heme sensors targeted to the nucleus and mitochondrial network (**Figure S1**), we found that HO-2 overexpression likewise depletes nuclear LH but does not affect mitochondrial matrix LH (**Figure 4d**).

**Figure 4.**
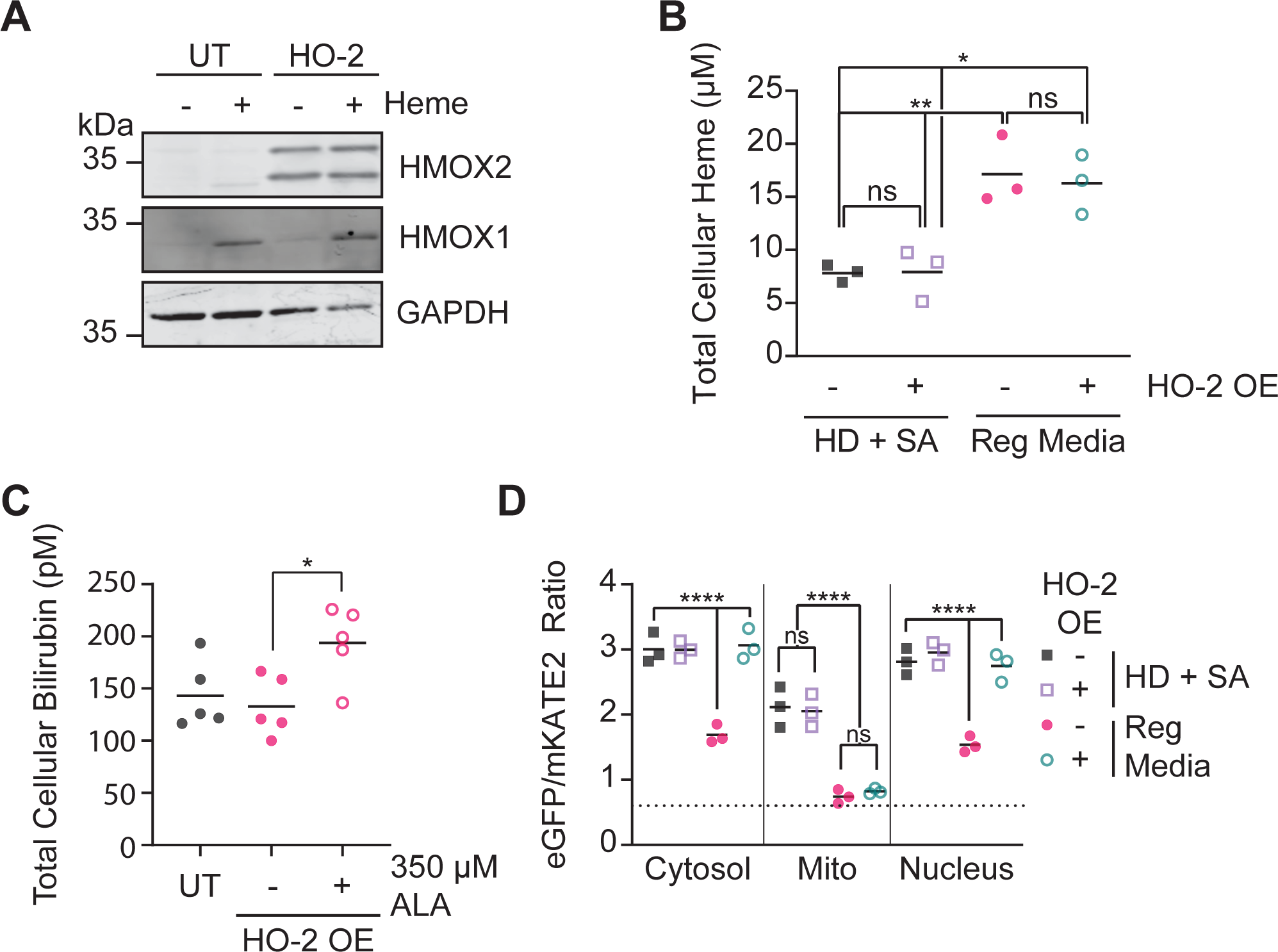
HO-2 overexpression depletes cytosolic and nuclear labile heme but not total heme, bilirubin levels, or mitochondrial labile heme. (**a**) Representative immunoblot of HO-1 (HMOX1), HO-2 (HMOX2), and GAPDH expression in untransfected (UT) or HO-2 overexpressing HEK293 cells treated with or without 10 µM hemin chloride. (**b**) Measurements of total heme in triplicate untransfected (-) or HO-2 overexpressing (OE) (+) HEK293 cultures grown in HD + SA or regular media. (**c**) Measurements of total bilirubin in triplicate untransfected (-) or HO-2 overexpressing (OE) (+) HEK293 cultures grown in regular media with or without 350 µM ALA. (**d**) Median cytosolic, nuclear or mitochondrial (mito)-targeted HS1 eGFP/mKATE2 fluorescence ratio values derived from flow cytometry experiments from triplicate untranfected (-) or HO-2 overexpressing (OE) (+) HEK293 cells grown in HD + SA or regular media. Representative violin plots of eGFP/mKATE2 fluorescence ratio distributions from single cell analysis of HEK293 cultures are shown in **Figure S8c**. The statistical significance is indicated by asterisks using one- way ANOVA for multiple comparisons using Tukey’s range test. *p<0.05, **p<0.01,***p <0.001, ****p<0.0001, ns = not significant.

We next sought to determine which features of the HO-2 polypeptide are required for its ability to deplete LH when overexpressed (**Figure 5a**). Mutation of the active site heme-binding histidine residue to alanine (H45A) results in a HO-2 mutant that is catalytically inactive, but retains the ability to weakly accommodate heme (64, 82). Overexpression of HO-2 H45A depletes LH, just as WT HO-2 (**Figure 5b**). However, consistent with the diminution of the HO-2 H45A affinity for heme, there is more LH in cells overexpressing the H45A mutant compared to WT HO-2 in regular media; the HS1 heme occupancies in cells overexpressing WT HO-2 and the H45A mutant are 6% and 20%, respectively.

**Figure 5.**
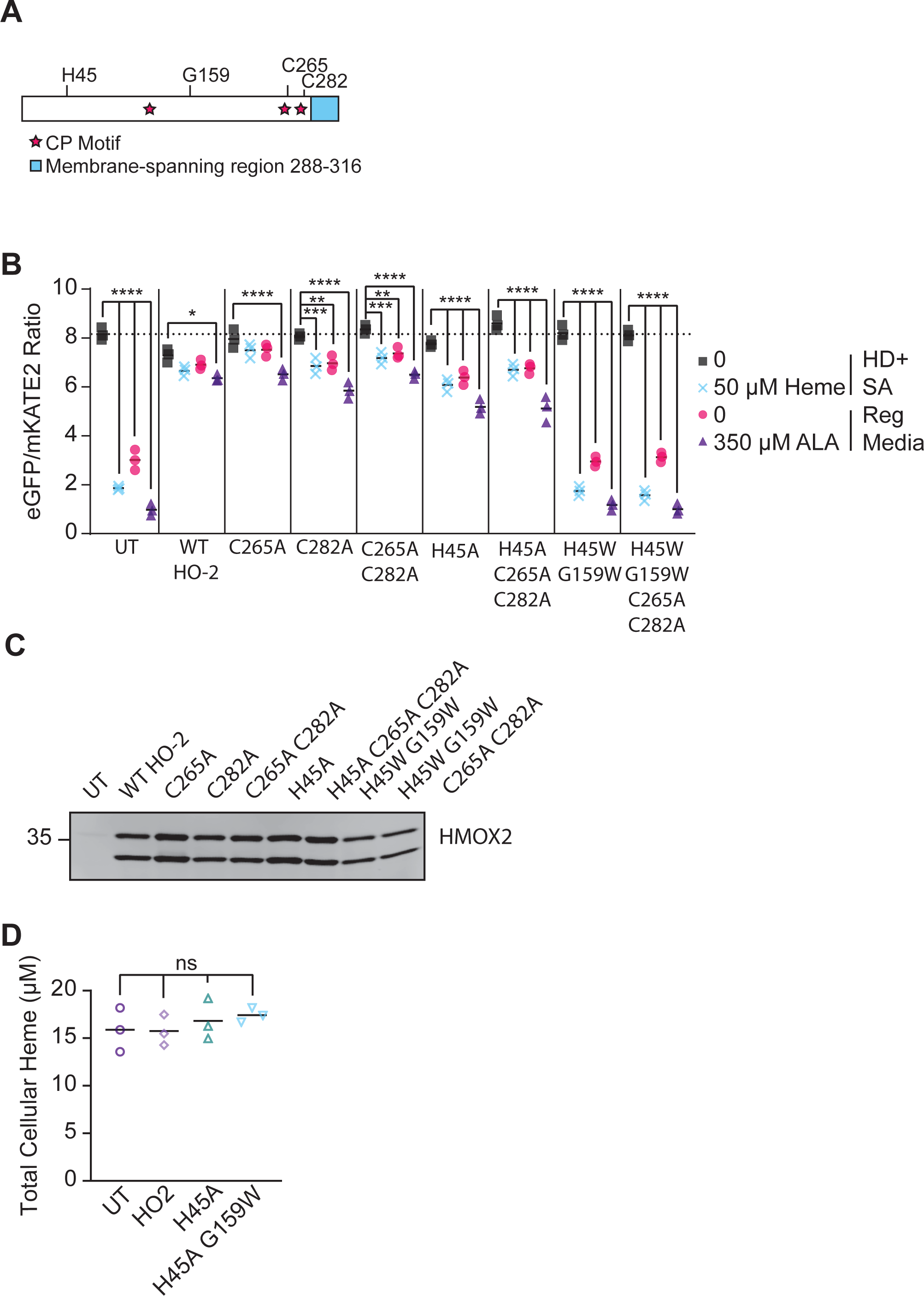
Heme sequestration and buffering requires the HO-2 heme binding pocket but not catalytic activity, heme regulatory motifs (HRMs), or ER membrane-tethering. (**a**) Schematic of HO-2 and residues of interest, including heme binding ligand H45, a residue that accommodates heme binding within the heme binding pocket, G159, Cys residues within Cys-Pro (CP; stars) dipeptides in the HRMs, C265 and C282, and the ER membrane-spanning region spanning residues 288-316 (blue). (**b**) Median cytosolic HS1 eGFP/mKATE2 fluorescence ratio values derived from flow cytometry experiments from triplicate HEK293 cells that were untransfected (UT) or overexpressing WT or the indicated mutant HO-2 alleles. Cells were cultured in HD + SA, treated with or without 50 µM hemin chloride, or regular media, treated with or without 350 µM ALA. Representative violin plots of eGFP/mKATE2 fluorescence ratio distributions from single cell analysis of HEK293 cultures are shown in **Figure S9**. (**c**) Representative immunoblot demonstrating overexpression of the indicated HO-2 (HMOX2) alleles relative to untransfected (UT) cells. (**d**) Measurements of total heme in triplicate cultures of HEK293 cells that were untransfected (UT) or overexpressing WT or mutant HO-2 grown in regular media. The statistical significance is indicated by asterisks using one-way ANOVA for multiple comparisons using Tukey’s range test. *p<0.05, **p<0.01,***p <0.001, ****p<0.0001, ns = not significant.

HO-2 also contains heme regulatory motifs (HRMs), which bind heme through cysteine within a cysteine-proline (CP) sequence and often a distal histidine (83, 84). In most proteins, the HRMs regulate protein degradation; however, in HO-2, the HRMs affect protein dynamics (39) and in transfer and loading of heme to the catalytic center (85), but does not otherwise affect protein stability or activity (64). We find that mutation of one or both C-terminal HRMs, C265A and/or C282A, do not affect the ability of overexpressed HO-2 to reduce LH (**Figure 5b**).

Lastly, we tested if mutation of the heme binding pocket affected LH. A H45W/G159W HO-2 mutant cannot bind heme (85) and is unable to deplete labile heme when over-expressed (**Figure 5b**). Importantly, the HO-2 mutations tested still resulted in high levels of steady-state expression of HO-2 (**Figure 5c**). Only the H45W/G159W mutant exhibited a decrease in protein expression compared to the other HO-2 variants, consistent with a prior study (64). However, the ∼50% decrease in expression of the H45W/G159W mutant is not the reason why this variant cannot deplete LH. If H45W/G159W HO-2 could deplete labile heme as the other HO-2 variants, but was expressed at half the level, then it would still diminish LH, but to a lower degree relative to the other HO-2 variants. Also of note, localization to the ER (**Figure S2**) and total heme levels (**Figure 5d**) are unaffected by the HO-2 mutations tested (64). Parenthetically, the ER-localization of HO-2 is not required for heme sequestration since a mutant lacking the ER-tethering C-terminal tail still depletes LH (**Figure S3**). Altogether, our data indicate that HO-2 overexpression depletes labile heme due to its ability to bind and sequester heme, not because of its ability to catalytically degrade heme.

### Labile heme is largely oxidized in HEK293 cells

Since HS1, which is ∼50% heme occupied in HEK293 cells, can bind both oxidation states of heme tightly, with *K*_DIII_ and *K*_*D*_^II^ values of 3 nM and 1 pM, respectively, the oxidation state of LH was unclear. The corresponding buffered free heme concentrations would span a range between 1 pM and 5 nM if it were fully reduced or oxidized, respectively (**Figure 2e**). Since HO-2 selectively binds ferric over ferrous heme, with a *K*_DIII_ = 3.6 nM (77) and *K*_*D*_^II^ = 320 nM (86), the observed changes in HS1 heme occupancy upon depletion or overexpression of HO-2 is consistent with LH being largely oxidized. If LH were reduced, changes in HO-2 expression over the concentration span observed in our work would not perturb HS1 heme occupancy due to the large difference in ferrous heme binding affinities, *K*_DHS1_ = 1 pM and *K*_DHO-2_ = 320 nM.

To graphically illustrate this point, HS1 heme occupancy as a function of HO-2 expression (**Figure 6**) was modeled using the equilibria depicted in **Equation 3** and described in greater detail in **Supplementary Information**.

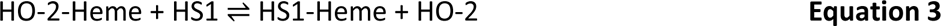

**Figure 6.**
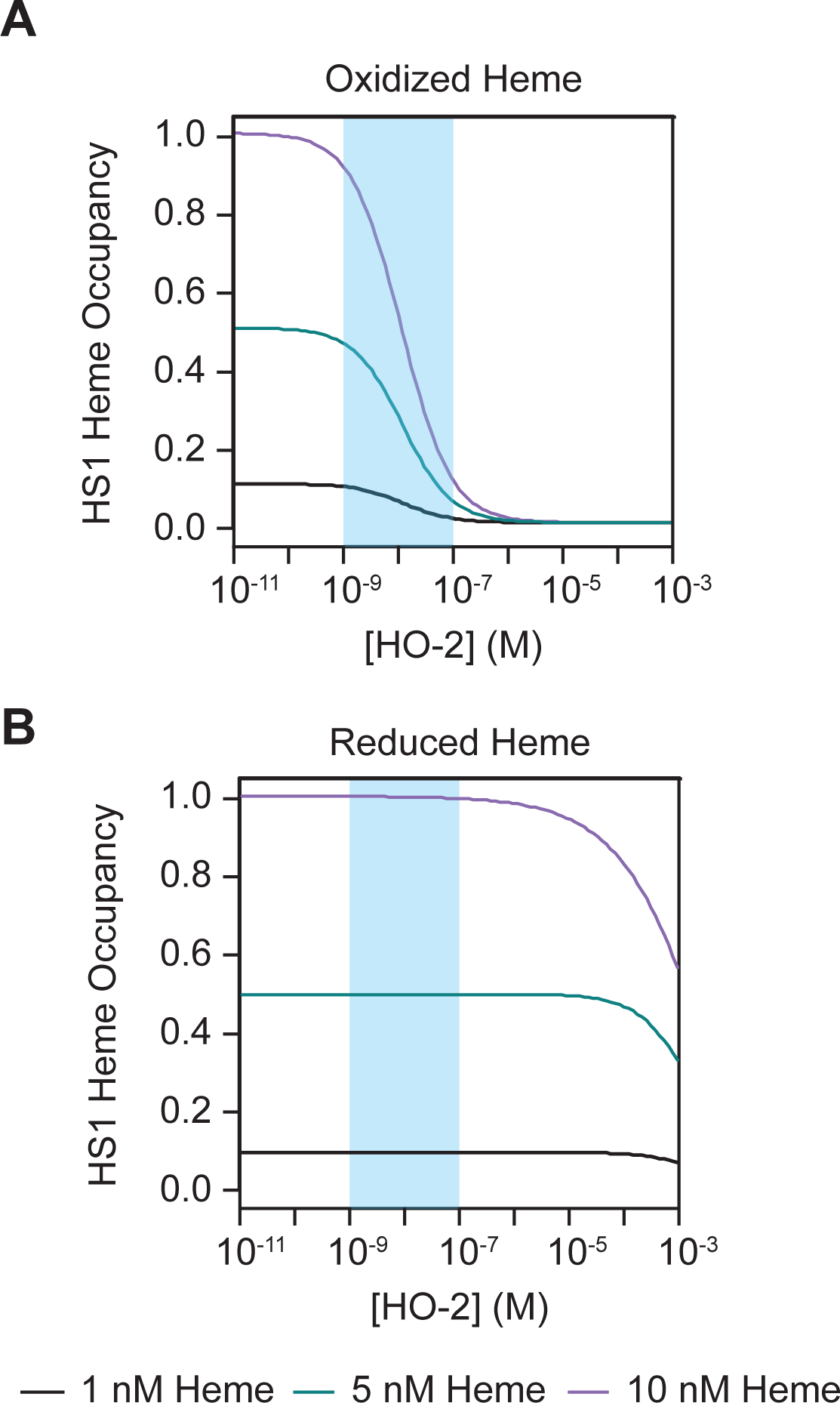
The effects of HO-2 depletion or overexpression on HS1 heme occupancy are consistent with labile heme being oxidized. (**a**) Simulation of HS1 heme occupancy as a function of HO-2 expression assuming the equilibrium model depicted in **Equation 3**, with [HS1] = 10 nM, and heme being oxidized and present at 1 nM (black), 5 nM (blue), or 10 nM (grey). (**b**) Simulation of HS1 heme occupancy as a function of HO-2 expression assuming the equilibrium model depicted in **Equation 3**, with [HS1] = 10 nM, and heme being reduced and present at 1 nM (black), 5 nM (blue), or 10 nM (grey). See text for details and **Supporting Information** for derivations of the equilibrium model. See also **Figure S4**, which depict simulations for 100 nM and 1 µM [HS1] and 0.1, 0.5, and 1.0 equivalents of heme relative to [HS1] in oxidized or reduced states.

Parameters of the model include the competition constant, *K*_Comp_, which is defined as the ratio of *K*_DHS1_ to *K*_DHO-2_, and the concentrations of HS1 and heme. *K*_Comp_ was fixed as either 0.8 for ferric heme (*K*_DHS1_ = 3.0 nM and *K*_DHO-2_ = 3.6 nM) or 3.3 x 10_-6_ for ferrous heme (*K*_DHS1_ = 1 pM and *K*_DHO-2_ = 320 nM). The HS1 heme occupancy was simulated for a variety of concentrations of sensor and heme **(Figure 6** and **Figure S4**). The HO-2 concentration in cells is 10 nM (**Figure S5**), with overexpression resulting in a ∼10-fold increase in expression (64) (**Figure 4a**) and silencing resulting in ∼80% reduction in expression (**Figure 3a**). The concentration of HS1 in cells is on the order of ∼10 nM (63, 67). If LH was oxidized, alterations in HO-2 expression between 1 - 100 nM would be expected to alter HS1 heme occupancy, as was observed (**Figure 6a**). On the other hand, if LH were reduced, HO-2 concentrations would have to approach mM levels to alter HS1 heme loading (**Figure 6b**). Taken together, the change in HS1 heme occupancy due to HO-2 expression is consistent with LH being oxidized. If one assumed HS1 were sensing free heme, it would translate to a buffered free heme concentration on the order of 5 nM in HEK293 cells. This conclusion supports our findings that heme may be too limiting for HO-2 (*K*_M_ = 400 nM) to be active in HEK293 cells and explains why bilirubin levels are not altered in response to HO-2 depletion or overexpression.

## Discussion

Heme oxygenases are arguably the best characterized systems for degrading and detoxifying excess heme (26–31). Mammals encode two HO isoforms, an inducible HO-1 and a constitutive HO-2 (32–37). The current conceptual paradigm for dual HOs is that HO-2 functions to degrade a baseline level of heme under homeostatic conditions and inducible HO-1 provides additional protection against heme toxicity in response to various stressors, including excess heme, oxidative insults, and exposure to various xenobiotics (57–61). However, prior to this work, it was unclear if endogenous heme was sufficiently available as a substrate for HO-2 (78).

*A priori*, in conceptualizing cellular heme, one can consider total heme to be the sum of the contributions from heme bound to exchange inert and labile heme complexes and unbound free heme. Therefore, heme availability to HOs is dependent upon its access to certain labile heme complexes that can exchange with HO or free heme. Degrading heme from exchange inert heme complexes requires a mechanism for heme dissociation, likely involving proteasomal or lysosomal protein degradation and ultimately release of heme into exchange labile heme complexes or an unbound free state. Given the large complement and concentration of biomolecules that could associate with heme, including proteins, nucleic acids, lipids, and an assortment of small molecules, free heme is buffered to very low concentrations.

The recent development of heme sensors from a number of labs has enabled measurements of chelatable heme. Strictly speaking, these sensors measure exchangeable heme *available* to the sensor, not necessarily *free* heme. Thus, sensor heme occupancy is the best parameter to report when interpreting changes in heme availability. On the other hand, if one assumes cellular chelatable heme is unbound, sensor heme occupancy values can be converted to a buffered free heme concentration. Using a number of different heme reporters across various eukaryotic cell lines, from yeast to a number of non-erythroid human cell lines, buffered free heme concentrations have been estimated to be between 5 and 600 nM (63,68,73,75). Given this ∼100-fold span in free heme, which is also a proxy for exchange labile heme, it was not known how active HO-2 would be. Moreover, since HO-2 binds ferric heme in its resting state (41–44), it was unclear what the oxidation state of the LH pool is or if HO-2 could even equilibrate with it.

Herein, on the basis of the changes in HS1 heme occupancy and the lack of change in total heme and bilirubin levels in response to HO-2 depletion and over-expression in HEK293 cells, we found that LH is largely oxidized and too limiting to support a prominent role for HO-2 in heme degradation. If assuming HS1 and HO-2 bind free heme, we can estimate a buffered free heme concentration on the order of 5 nM, which is identical to that reported recently using a peroxidase-based heme reporter in HEK293 cells (69), making heme too limiting to support heme degradation by HO-2 (*K*_MHeme_ = 400 nM) (37). If assuming HS1 and HO-2 access heme via exchange reactions with certain labile heme complexes, heme exchange with HO-2 is likewise too limiting to support heme degradation. In this case, the HO-2 free heme *K*_M_ value is physiologically irrelevant, and a screen for heme-protein complexes that can exchange with HO- 2 and knowledge of their *K*_M_ values with HO-2 is needed to better understand heme substrate availability and HO-2 activity *in vivo*. Regardless of what formalism one uses to contemplate heme availability to HO, *i.e.* free heme vs. exchangeable heme, we found that HO-2 plays a role in regulating heme bioavailability through its ability to bind and buffer heme, not by degrading it as is typically assumed. Our results have a number of important physiological implications for heme availability, signaling, and heme oxygenase function.

The oxidation state of LH in eukaryotic cells has been a long-standing unresolved mystery. LH was first characterized *in situ* in live human cell lines with genetically encoded FRET-based heme sensors (68). In these seminal studies, only the ferric heme affinities of the sensors were determined and estimates of buffered free heme concentrations based on sensor heme occupancy were by default assumed to be ferric. These studies found free heme to be ∼20 nM in a variety of non-erythroid human cell lines, including HEK293 cells. Studies using heme-peroxidase activity-based reporters, which binds ferric heme in its resting state, found free heme to be as high as ∼400 nM in HEK293 cell lysates (73). These estimates may be higher since heme measurements were done in disrupted cell extracts and cell lysis may have resulted in heme dissociation from certain heme binding sites. Interestingly, unlike in HEK293 cells, in *Saccharomyces cerevisiae*, heme occupancies for the high affinity heme sensor, HS1, was found to be 100% and the lower affinity sensor, HS1-M7A, was 30-50% heme loaded. At the time of these studies, it was assumed that LH was largely reduced on the basis of the highly reducing environment of the cell, *E*_M_ = -330 mV (87), which is dictated by the glutathione redox buffer, and the reduction potential of monomeric aqueous heme, *E*_M_ = -50 mV (63,88,89). The heme occupancy of HS1-M7A coupled with its *K*_*D*_^II^ value of 25 nM led to estimates of free heme being ∼25 nM (if LH was primarily reduced) in yeast. Altogether, prior to this study, there were no methods to parse LH oxidation state.

In the current report, we exploited HO-2 selectivity for ferric heme over ferrous heme (86) and perturbations to its expression by silencing or over-expression to infer that LH is largely oxidized (**Figure 6**). These findings suggest that the glutathione redox buffer does not equilibrate with LH to keep it reduced and supports prior studies that found glutathione binds oxidized heme too weakly (*K*_DIII_ = 20 μM) to represent a physiologically relevant buffering factor (90). LH being oxidized is consistent with the observation that DNA and RNA guanine quadruplexes, which have nM affinities for ferric heme, also appear to bind and regulate heme availability in human cell lines (67, 71). Moreover, LH being oxidized is also consistent with previously reported roles of oxidized heme in signaling. For instance, ferric, but not ferrous, heme is required for RNA-binding protein DGCR8 for primary microRNA processing (91). The nuclear receptor Rev-Erbβ can sense and bind ferric heme, which can then act as a sensor for CO or NO by coupling gas binding to heme reduction (92). In addition, ferric heme was demonstrated to regulate ATP dependent potassium channels (93).

Although LH is largely oxidized in HEK293 cells, it may vary between different cell types and be highly dynamic and responsive to various stimuli. The Fe(III)/Fe(II) redox couple is linked to a number of factors, including access to certain cellular reductants or oxidants (94), ligand binding to the heme iron center, *e.g.* CO or NO (92), and allosteric protein conformational changes (95). Exactly how these factors conspire to affect steady-state LH oxidation state, its dynamics, and role in metabolism and physiology is not known and may be elucidated by the future development of oxidation-state specific heme sensors or chelators.

Our results suggesting that endogenous LH or free heme is too limiting in some cell types to support HO activity has implications for the function of HO-2, HO-1, and emerging roles for heme catabolites, *e.g.* CO, biliverdin, and bilirubin in physiology. First, our results explain why overexpression of catalytically inactive HO-2 alleles are cytoprotective against peroxide stress (76). It likely does so by sequestering heme, presumably to prevent heme-dependent peroxidase or Fenton reactions that are damaging to cells. By extension, we suggest that non-heme stressors, *e.g.* ROS, metals, and various xenobiotics, that induce HO-1 expression may result in cytoprotection due to HO-1 mediated heme sequestration because endogenous LH or free heme may be too limiting in many cell types. The heme sequestration mechanism for heme detoxification may also occur if electron delivery to the HOs via CPR is limiting. During heme stress, and provided that there is sufficient CPR and NADPH, inducible HO-1 and constitutive HO-2 are expected to enzymatically degrade and detoxify excess heme, with their relative contributions being dictated by their respective expression levels.

The ratio of HO-2 to HO-1 protein expression is highly variable across different cell lines and tissues, spanning 5000 (frontal cortex) to 0.03 (spleen) (**Figure S6**) (96). Moreover, HO-1 can be induced by as much as 100-fold in response to various stressors (97). Thus, cells occupy a position along an expansive continuum between their reliance on HO-1 and HO-2. The cell line used in this study, HEK293 cells, has a HO-2 to HO-1 ratio of 2 (96, 98), explaining why silencing of HO-2 still results in measurable bilirubin levels. In most cell types, it is not known if free or labile heme is too limiting to support a significant amount of HO activity in the absence of excess heme stress, as it is in HEK293 cells, raising the intriguing possibility that HO-2, and possibly HO-1, have other roles in heme homeostasis beyond heme degradation.

Consistent with a potential role for HO-2 in controlling access to heme, it was recently demonstrated that HO-2 is degraded by the lysosomal pathway during heme deficiency (64), presumably to increase heme availability. A role for HO-2 in regulating heme availability needs to be explored further and could represent a mechanism for heme sparing during periods of heme limitation. On the other hand, there may be certain cell types in which endogenous LH or free heme is much higher than HEK293 cells, *e.g.* cells of the brain and testes, where HO-2 is most highly expressed. Indeed, prior work suggests that, in the brain, HO-2-derived heme catabolites can provide protection against oxidative stress and other injuries (31,99,100). Our future studies will involve profiling LH in multiple cell types with varying levels of HO-2 and HO- 1 expression to better assess HO function with respect to its heme degrading and buffering functions.

CO, biliverdin, and bilirubin, products of heme degradation, have emerged as key metabolites with distinct physiological roles (32,47–55). The only source of these metabolites is from heme degradation via the heme oxygenase system. Our studies in HEK293 cells finding that LH is too limiting to support catalytic heme degradation via the heme oxygenase system indicates that heme catabolite signaling in many cell types must be coupled to substantially increased heme synthesis, redistribution of LH, or influx of heme. In *Saccharomyces cerevisiae*, NO signaling (63) and heavy metal stress (62) was found to increase LH, suggesting that other physiological inputs may augment LH to support heme catabolite production. In addition, given the identification of mammalian heme importers and exporters, cells may communicate via heme, ultimately transducing heme signals to heme catabolites via the HOs to regulate physiology and metabolism.

Most interestingly, we found that HO-2 overexpression limits heme availability in the cytosol and nucleus, but not the mitochondria (**Figure 4d**). These data indicate that the mitochondrial LH pool is insulated from the generation of a cytosolic heme sink. In contrast, if mitochondrial LH is populated with exogenously derived heme, by supplementing cells cultured in HD + SA media with 6 μM hemin chloride, overexpression of HO-2 does reduce mitochondrial LH (**Figure S7**). Together, these data suggest that mitochondrially produced heme flows in only one direction, from the inside out, and not vice versa. In contrast, exogenously supplied heme can gain access to the mitochondria, but in a manner that is impacted by a cytosolic heme sink. These data are consistent with prior findings that exogenously supplied and endogenously synthesized heme are trafficked through separate pathways (73). Indeed, we found herein that although 350 µM ALA and 50 µM heme supplementation led to similar total intracellular heme levels, only the latter was able to populate the lower affinity HS1-M7A heme sensor (**Figure 2c** and **d**), further highlighting the differences in trafficking and bioavailability between exogenous and endogenous heme sources.

Altogether, our studies demonstrate that HO-2 may not be an active heme catabolizing enzyme in certain cell types due to heme substrate being limiting, raising the prospect that it may have alternative roles in regulating heme homeostasis. The identification of a heme buffering function for HO-2 in regulating heme availability places it with other factors, such as glyceraldehyde phosphate dehydrogenase (GAPDH) (63,70,101,102), rRNA and DNA guanine quadruplexes (67, 71), and mitochondrial-ER contacts sites (66), that affect intracellular heme availability and distribution. Future work is required to firmly establish the physiological context and consequences of HO-2 mediated regulation of heme availability independent of its role in heme degradation.

## Experimental Procedures

### Materials

Fetal bovine serum (FBS) (Catalog # 89510-188), Dulbecco’s Modified Eagle’s Medium (DMEM) (Catalog # L0102-0500), Trypsin 0.25% (Catalog # 45000-664), 5- aminolevulinic acid (ALA) (Catalog # BT143115-2G), ascorbic acid, cell culture grade DMSO, oxalic acid, Dulbecco’s Phosphate Buffered Saline (DPBS), and LiCOR Intercept Tris buffered saline (TBS) blocking buffer (Catalog # 103749-018) were purchased from VWR/Avantor. Opti-MEM (Catalog # 31-985-070) was purchased from Fischer Scientific. Lipofectamine LTX and PLUS reagent (Catalog # 15338100), Lipofectamine RNAiMax (Catalog # 13778075) and a mounting reagent with DAPI were purchased from Invitrogen. Succinyl acetone (Catalog # D1415) was purchased from Sigma Aldrich. Digitonin and hemin chloride were acquired from Calbiochem. The following primary and secondary antibodies were utilized: rabbit polyclonal heme oxygenase 2 antibody (Abcam ab90515); mouse monoclonal GAPDH antibody (Sigma 8795); rabbit polyclonal HO-1 antibody (Enzo Life Sciences BML-HC3001-0025); mouse monoclonal calnexin antibody (Invitrogen MA3-027); mouse monoclonal ubiquitin antibody (P4D1) (Santa Cruz Biotechnology sc-8017); anti-FLAG M2 magnetic bead (Sigma Aldrich M8823); goat anti-rabbit secondary antibody (Biotium 20064); and goat anti-mouse secondary antibody (Invitrogen SA5-35521).

### Plasmid constructs and mutagenesis

The previously described FLAG-tagged WT and mutant HO2 constructs were subcloned into the pcDNA3.1 plasmid and driven by the CMV promoter (64). For untargeted cytosolic expression of human codon optimized heme sensors, HS1, HS1- M7A, and HS1-M7A,H102A, the heme sensors were expressed using the pcDNA3.1 plasmid and driven by the CMV promoter as previously described (63, 67). For heme sensing experiments involving mitochondrial matrix and nuclear targeting of HS1, constructs were generated via Gateway Cloning (Invitrogen Life Technologies) by PCR amplifying the hHS1 constructs with primers containing 5’-attB1 (G GGG ACA AGT TTG TAC AAA AAA GCA GGC T) and 3’-attB2 (GGG GAC CAC TTT GTA CAA GAA AGC TGG GT) sequences. The purified PCR fragments were cloned into pDONR221 vector via BP recombination reaction. The resulting entry clones were subjected to LR recombination reaction with pEF5/FRT/V5-DEST vector to obtain the expression plasmids. The mitochondrial and nuclear targeting sequences utilized were from subunit VIII of human COXIV, MSVLTPLLLRGLTGSARRLPVPRAKIHSL, and a nuclear localization sequence, MDPKKKRKVDPKKKRKV (73). Sequences and plasmid maps are provided in **Supplementary Information**. All plasmids were confirmed by Sanger sequencing.

### Cell culture, growth conditions, plasmid transfections, and RNA interference

HEK293 cells used in this study were previously described and obtained from American Type Culture Collection (ATCC) (63, 67). HEK293 cells were grown in regular media - Dulbecco’s Modified Eagle’s Medium (DMEM), with 4.5 g/L Glucose and without L-Glutamine and Sodium Pyruvate, supplemented with 10% FBS (Heat Inactivated) in T75 flasks (Greiner) (Regular Media). Cells were split using .25% Trypsin.

To deplete heme, cells were cultured with 500 μM succinylacetone (SA) in heme deficient (HD) media - DMEM containing 10% heme depleted FBS - for 48-72 hours prior to harvesting. To increase heme, cells were cultured with 350 μM 5-aminolevulinic acid (ALA) or between 1 and 50 μM hemin chloride in regular or HD media for 20-24 hours. Heme-depleted FBS was generated by incubating it with 10 mM ascorbic acid in a 37°C shaking incubator at 200 RPM for 8 hours. Loss of heme was monitored by measuring the decrease in Soret band absorbance at 405 nm; typically the initial Abs_405nm_ was 1.0 and the final Abs_405nm_ was 0.5. 80 mLs of ascorbate treated FBS was then dialyzed against 3 L of 1X PBS, pH 7.4, three times for 24 hours each at 4 °C, using a 2000 MWCO membrane (Spectra/Por 132625). Dialyzed FBS was then filter sterilized with a 0.2 μm polyethersulfone filter (VWR 28145-501) and syringe (VWR 53548-010).

For heme sensor transfections, 1/24 of the HEK293 cells from a confluent T75 flask were used to seed 2 mL cultures in each well of polystyrene coated sterile 6 well plates (Greiner). Once cells were 30% - 50% confluent, 1-2 days after seeding, the media was changed to 2 mLs of fresh regular or HD+SA media 30 - 60 minutes before transfection. For each well to be transfected, 100 µL of a plasmid transfection master mix was added. The master mix was prepared by mixing, in order, 600 µL OptiMEM (100 µL per well up to 600 µL), 2 µg of heme sensor plasmid, a volume of Lipofectamine Plus Reagent equal in µL to the µg of plasmid DNA used, and a volume of Lipofectamine LTX transfection reagent that is double the volume of the Lipofectamine Plus Reagent. Prior to adding the transfection mixture to each 6-well plate, the master mix was mixed by gentle pipetting and allowed to incubate at room temperature (25 °C) for 5 minutes. 48 hours after transfection, the media was changed to fresh regular or HD+SA media, and supplemented with the appropriate concentration of ALA or hemin chloride as indicated. Cells were then cultured for an additional 20-24 hours prior to harvesting for various analyses, including measurements of labile heme, total heme, and bilirubin, as well as immunoblotting.

If co-transfecting the heme sensors with the HO2 plasmids, the same procedure was followed as above, with the exception that 2 µg of the HO-2 plasmid DNA was added at the same time as the heme sensor plasmid, and twice the volumes of the Lipofectamine Plus and LTX reagents were used.

For cytosolic, mitochondrial, or nuclear targeted pEF-hHS1 in which HO-2 plasmids were not co-transfected, PolyJET was used in plasmid transfections. The procedure is identical to that described above, except that 100 μL of a PolyJET transfection reagent master mix was added to each well in a 6-well plate. Per well, the master mix consisted of 50 μL of DPBS with 1-2 μg of plasmid DNA and 50μL of DPBS with 4-8 μL of PolyJET, and was incubated at room temperature for 10-15 minutes prior to adding to the cells. 24-40 hours post transfection, the media was changed and cells were then cultured for an additional 20-24 hours with ALA or hemin chloride as indicated prior to harvesting for labile heme and total heme measurements.

In order to silence HO-2 expression in HEK293 cells, we used siRNA against HO-2. HEK293 cells were plated in polystyrene coated sterile 6 well plates (Greiner) and cultured in regular media as described above. At 10-30% confluency, cells were transfected with siHMOX2 (Dharmacon L-009630-00-0005) or non-targeting pool (Dharmacon D-001810-10-05) siRNA using Lipofectamine RNAi Max (Thermofisher). Briefly, for each well to be transfected, 9 μL of Lipofectamine RNAiMax was added to 150 μL of OptiMEM, and 3 μL of 10 μM siRNA to 150 μL OptiMEM. The two solutions were mixed 1:1 and allowed to incubate at room temperature for 5 minutes. After incubation, 250 μL of the transfection solution was added to the 2 mL cultures. After 24 hours, the HO-2 silencing protocol was repeated and the cells were cultured for an additional 48 hours. After 72 hours of control or siRNA treatment, cells were harvested for analysis.

In order to transfect the heme sensor plasmids into HO-2 silenced cells, the pEF5-hHS1 plasmid was transfected using the Lipofectamine LTX reagents as described above after 72 hours of control or siRNA treatment. After transfecting the heme sensor, cells were cultured for an additional 72 hours in either HD+SA or regular media. In order to induce heme excess, cells cultured in regular media were treated with 350 μM ALA for 24 hours prior to harvesting.

### Heme sensor calibration

In order to determine sensor heme occupancy, *R*_min_ and *R*_max_ was determined by depleting heme from cells or saturating the sensor with heme, respectively (**Equation 1**). Heme depletion was conducted by growing parallel cultures in HD + SA media for 3 days, as described above. To saturate the sensor with heme, cells from a well of a 6-well dish were resuspended in 1 mL of “sensor calibration buffer”, transferred to a microfuge tube, and incubated in a 30 °C water bath for 30 minutes. Following this, cells were pelleted at 400 x g for 4 minutes at 4 °C, the media aspirated, the cell pellet washed with 1 mL PBS at room temperature once, and finally resuspended in 500 μL PBS containing 1mM ascorbate before being analyzed by flow cytometry. The sensor calibration buffer consisted of 1 mL of serum-free DMEM (pre-warmed at 37 °C), 40 μM digitonin (from a 1 mg/mL stock solution in PBS), 100 μM hemin chloride (from a 10 mg/mL DMSO stock) and 1 mM ascorbate (from a 100 mM stock solution in PBS). The sensor calibration buffer saturates cytosolic and nuclear hHS1 and partially saturates mitochondrial hHS1. An alternative method to saturate hHS1 in all compartments tested, including the mitochondria, involved culturing cells in 350 µM ALA for 24 hours prior to fluorescence analysis.

### Flow Cytometry

HEK293 cells were transfected and cultured as described above in 6 well polystyrene coated plates. Prior to analysis, cells were washed and resuspended in 1 mL of 1X PBS and transferred to a 1.5 mL microfuge tube and pelleted at 400 rcf 4°C for 4 min. Supernatant was decanted and cells were resuspended in 500 μL 1X PBS. Cell suspension was filtered through 35 μm nylon filter cap on 12x75 mm round bottom tubes (VWR/Falcon 21008- 948). Flow cytometric measurements were performed using a BD LSR Fortessa, BD LSR II, or BD FACS Aria Illu flow cytometers equipped with an argon laser (ex 488 nm) and yellow-green laser (ex 561 nm). EGFP was excited using the argon laser and was measured using a 530/30 nm bandpass filter, mKATE2 was excited using the yellow- green laser and was measured using a 610/20 nm bandpass filter. Data evaluation was conducted using FlowJo v10.4.2 software. Single cells were gated by size (FSC versus SSC) and only mKATE2 positive cells that had median mKATE2 fluorescence were selected for ratiometric analysis, which typically corresponded to ∼5000 cells.

### Bilirubin measurements

HEK293 cells were transfected and cultured as described above in 6 well polystyrene coated plates. Prior to analysis, cells were washed and resuspended in 1 mL of 1X PBS and transferred to a 1.5 mL microfuge tube and pelleted at 400 rcf 4°C for 4 min. Supernatant was decanted and cells were resuspended in 1 mL 1X PBS, and counted using a TC20 Automated Cell Counter from BioRad. 1x10^6^ cells were pelleted and resuspended in Lysis Buffer (10 mM NaPi 50 mM NaCl, 1% Triton X-100, 5 mM EDTA, 1 mM PMSF, 1x Protease Arrest (G-Bioscience 786-437)), then lysed by freeze-thawing using 3 cycles of 10 minutes at -80 °C and 15 minutes at RT. Lysates were clarified by centrifugation at 21.1xg in a table-top microfuge at 4°C for 10 minutes. 1% DMSO was added to the cell lysate to solubilize the bilirubin. Bilirubin was quantified using the Bilirubin Assay kit from Sigma-Aldrich exactly as described in the manufacturer’s protocol (MAK126). A standard curve was generated from bilirubin standards made up in cell lysis buffer with 1% DMSO. Bilirubin concentrations were normalized to cell number and a cellular concentration was determined by assuming a cell volume of 1.2 pL (103).

### Total heme measurements

Measurements of total heme were accomplished using a fluorometric assay designed to measure the fluorescence of protoporphyrin IX upon the release of iron from heme as previously described. Following the harvesting of HEK293 cells as described above, cells were counted using an automated TC20 cell counter (BioRad). At least 500,000 cells were resuspended in 400 µLs of 20 mM oxalic acid overnight at 4°C protected from light. Following the overnight incubation, an equal volume of 2 M warm oxalic acid was added to give a final oxalic acid concentration of 1M. The oxalic acid cell suspensions were split, with half the cell suspension transferred to a heat block set at 100 °C and heated for 30 minutes and the other half of the cell suspension kept at room temperature (∼25 °C) for 30 minutes. All suspensions were centrifuged for 3 minutes on a table-top microfuge at 21000 x g and the porphyrin fluorescence (ex: 400 nm, em: 620 nm) of 200 µL of each sample was recorded on a Synergy Mx multi-modal plate reader using black Greiner Bio-one flat bottom fluorescence plates. Heme concentrations were calculated from a standard curve prepared by diluting 2.5- 200 nM hemin chloride stock solutions in 0.1 M NaOH into oxalic acid solutions prepared the same way as for the cell samples. In order to calculate heme concentration, the fluorescence of the unboiled sample (taken to be the background level of protoporphyrin IX) is subtracted from the fluorescence of the boiled sample (taken to be the free porphyrin generated upon the release of heme iron). The cellular concentration of heme is determined by dividing the moles of heme determined in this fluorescence assay and dividing by the number of cells analyzed, giving moles of heme per cell, and then converting to a cellular concentration by dividing by the volume of a HEK293 cells, assumed to be 1.2 pL (103).

### Immunoblotting

Cells were harvested from a 6 well plate in 1X PBS and pelleted at 21100 x g at 4°C for 1 min and resuspended in lysis buffer (10 mM NaPi 50 mM NaCl, 1% Triton X-100, 5 mM EDTA, 1 mM PMSF, 1x Protease Arrest (G-Bioscience 786-437)). Cells were lysed by freeze-thawing using 3 cycles of 10 minutes at -80 °C and 15 minutes at RT. Lysates were clarified by centrifugation at 21.1xg in a table-top microfuge at 4°C for 10 minutes. Lysate protein was quantified by Bradford Assay. Typically, 15-30 µg of lysate protein was prepared in a final volume of 20 uL loading buffer, with 1x SDS loading dye in lysis buffer (5x SDS loading dye consists of 312.5 mM Tris-HCl, pH 6.8, 50% Glycerol, 346 mM SDS, 518 mM DTT, 7.5 mM Bromophenol blue). Samples were boiled at 100°C for 5 minutes in a heat block. Lysate samples were loaded onto a 14% Tris-Glycine SDS-PAGE gel (Invitrogen XP00140BOX) and electrophoresed at a constant current of 20 mA for 1.5 hours. Gels were transferred onto 0.2 um nitrocellulose membranes (Biotrace) overnight, but no longer than 2 days, using the XCell II Blot Module (Invitrogen) for a wet tank transfer. Membranes were blocked with Intercept TBS Blocking Buffer (LICOR 927-50000). HO-2 and HO-1 were probed with a 1:1,000 dilution of rabbit anti HO-2 antibody (Abcam 90515) or 1:1,000 rabbit anti HO-1 (Enzo Life Sciences BML- HC3001-0025) at 4°C overnight, respectively. GAPDH was probed with a 1:4,000 dilution of mouse anti GAPDH (Sigma 8795) for 1 hour at room temperature. Goat anti rabbit secondary antibody conjugated to a 700 nm fluorophore (Biotium 20064) or goat anti mouse secondary antibody conjugated to a 800 nm fluorophore (Invitrogen SA5-35521) was used at a dilution of 1:10,000 for 1 hour at room temperature. After application of primary and secondary antibodies, membranes were washed with 1X Tris Buffered Saline-Tween (TBS-T) (24.7 mM Tris-HCl pH 7.4 13.6 mM NaCl, 2.6 mM KCl 0.1% Tween 20) two times for 10 minutes at room temperature. Membranes were imaged using a LiCOR Odyssey Clx imager.

### Immunofluorescence microscopy (IF)

The experimental approach for IF are adapted from previous literature (64). Briefly, HEK293 cells were cultured on poly-lysine coated coverslips, transfected with plasmids expressing FLAG-tagged HO2 and its variants, washed with phosphate buffered saline (PBS) and fixed with 4% paraformaldehyde (w/v) before immunostaining. IF samples were permeabilized with 1x PBS containing 0.2% Triton X-100 (v/v) and stained with corresponding antibodies. Sample images were acquired on an Olympus IX81 microscope with a Yokogawa CSU-W1 spinning disk confocal scanner unit, an Olympus PlanApo 60x 1.42 NA objective, and a Hamamatsu ImagEMX2-1K EM-CCD digital camera.

### Supporting Information

This article contains supporting information.

## Acknowledgments

We acknowledge support from the core facilities at the Parker H. Petit Institute for Bioengineering and Bioscience at the Georgia Institute of Technology for expert advice and use of equipment.

---

**Figure S1.**
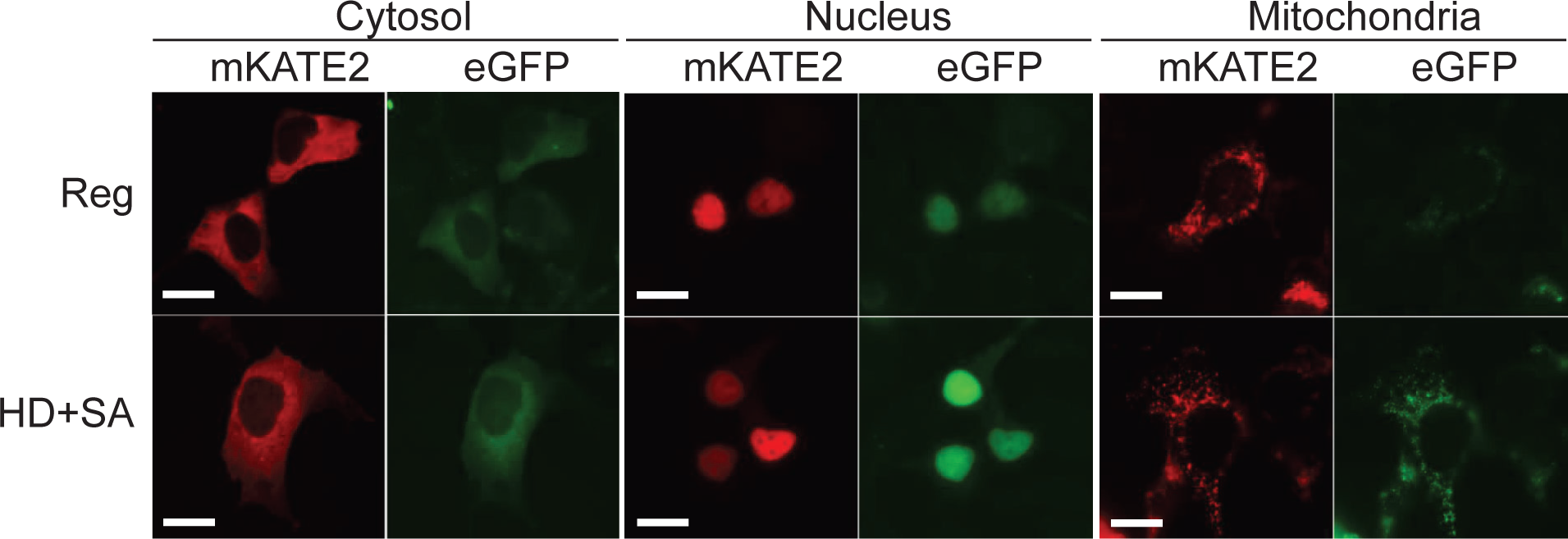

**Figure S2.**
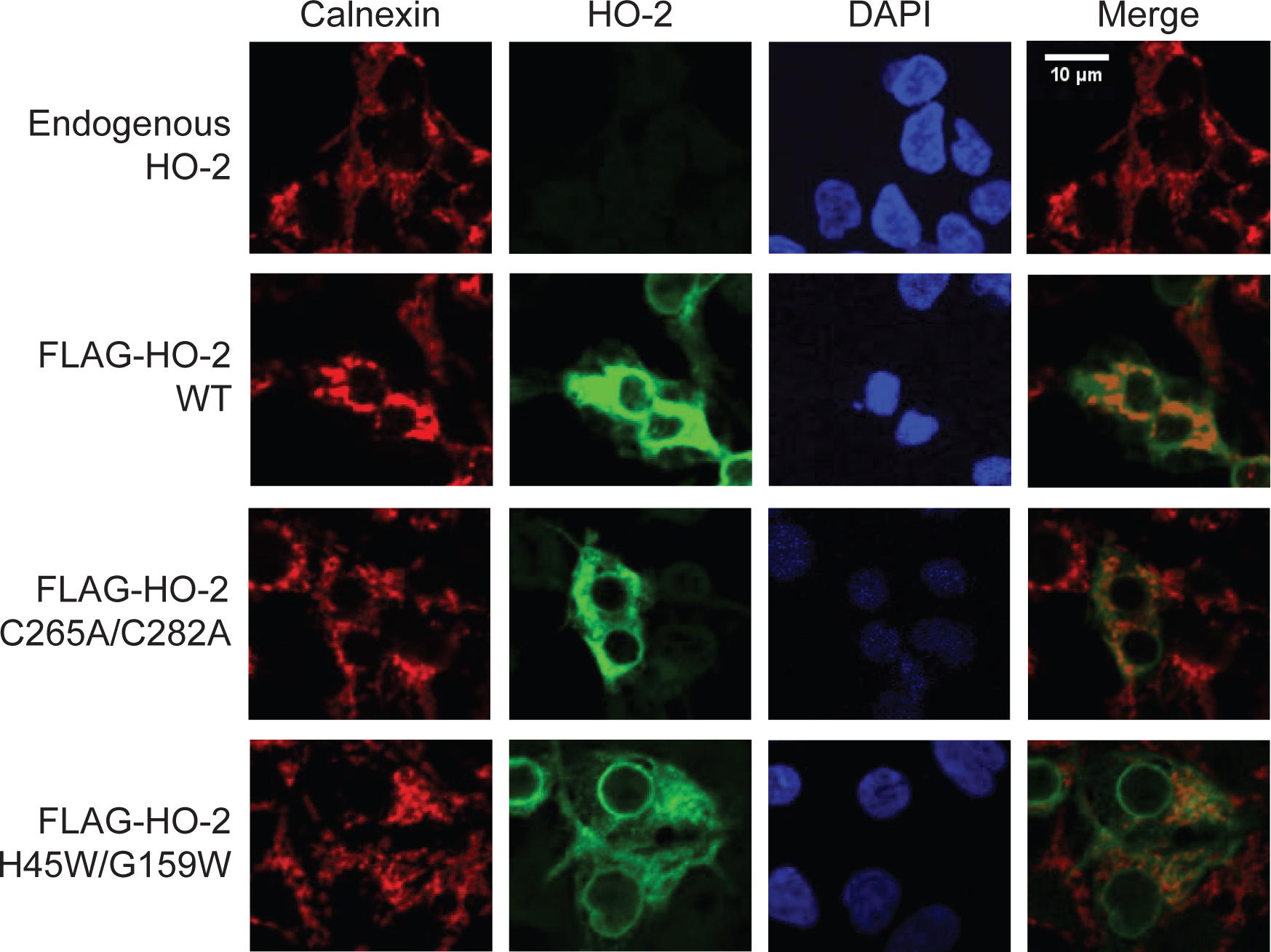

**Figure S3.**
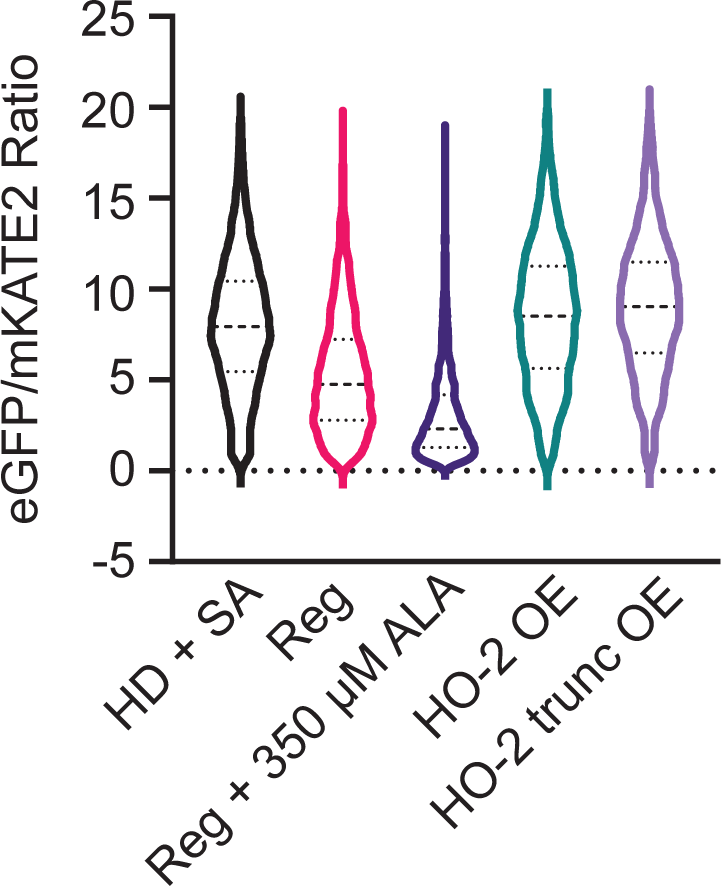

**Figure S4.**
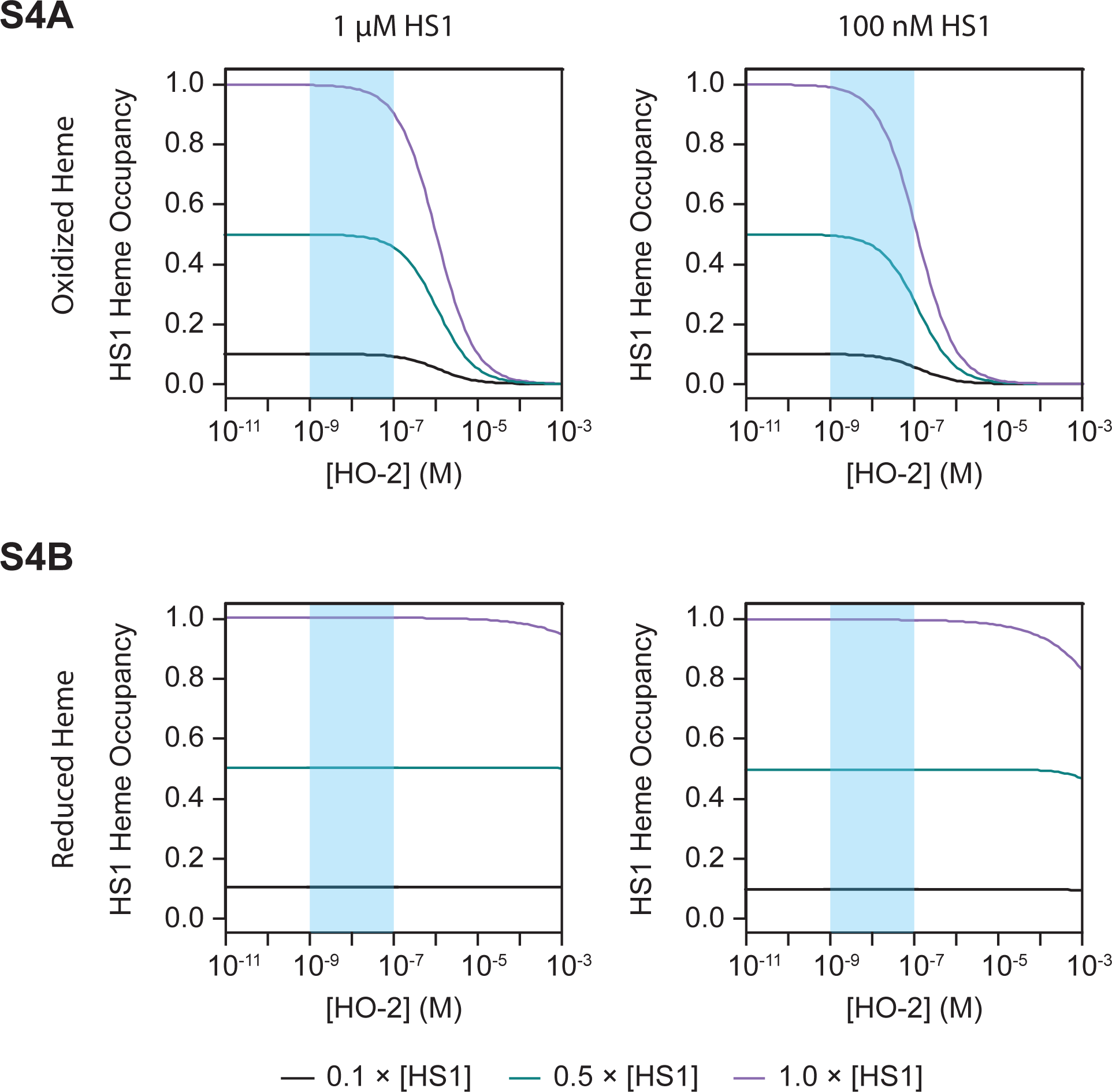

**Figure S5.**
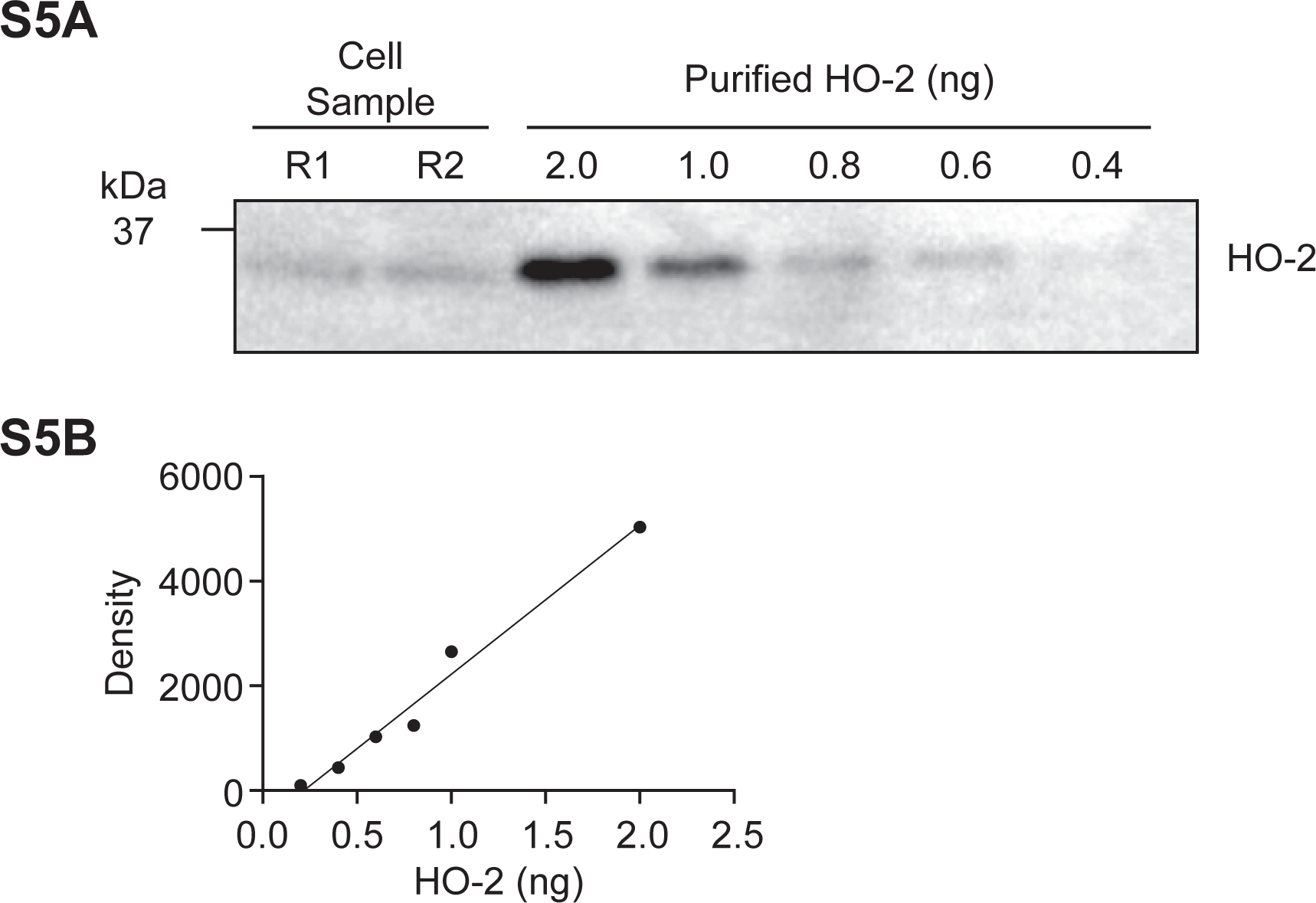

**Figure S6.**
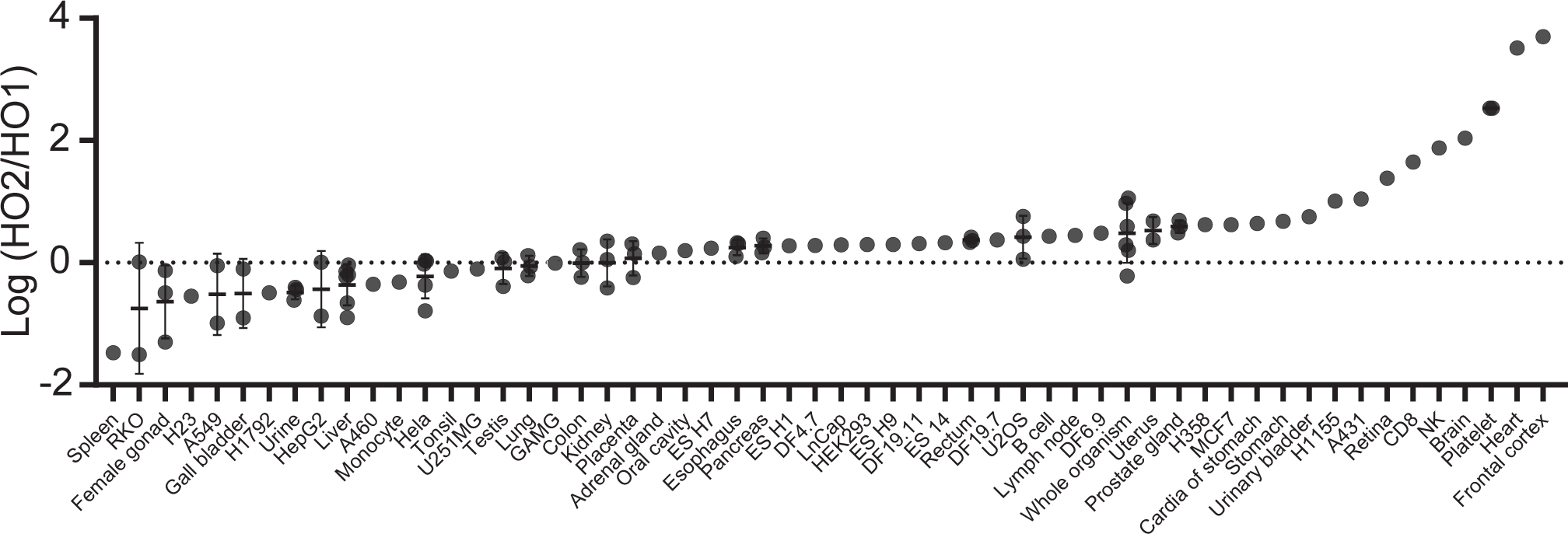

**Figure S7.**
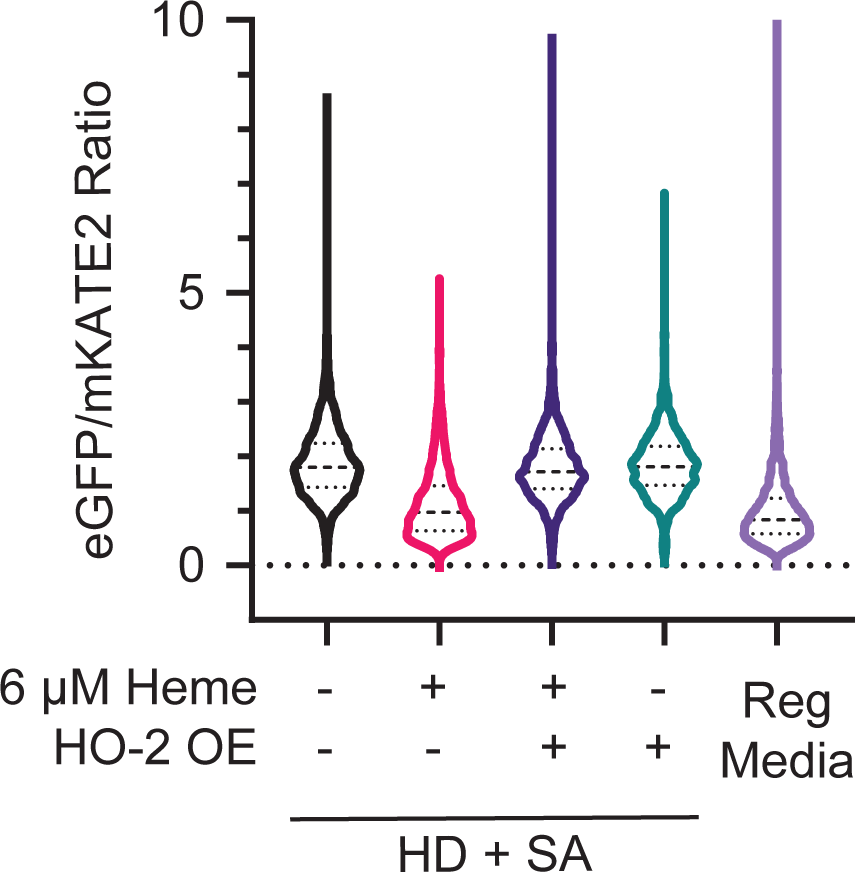

**Figure S8.**
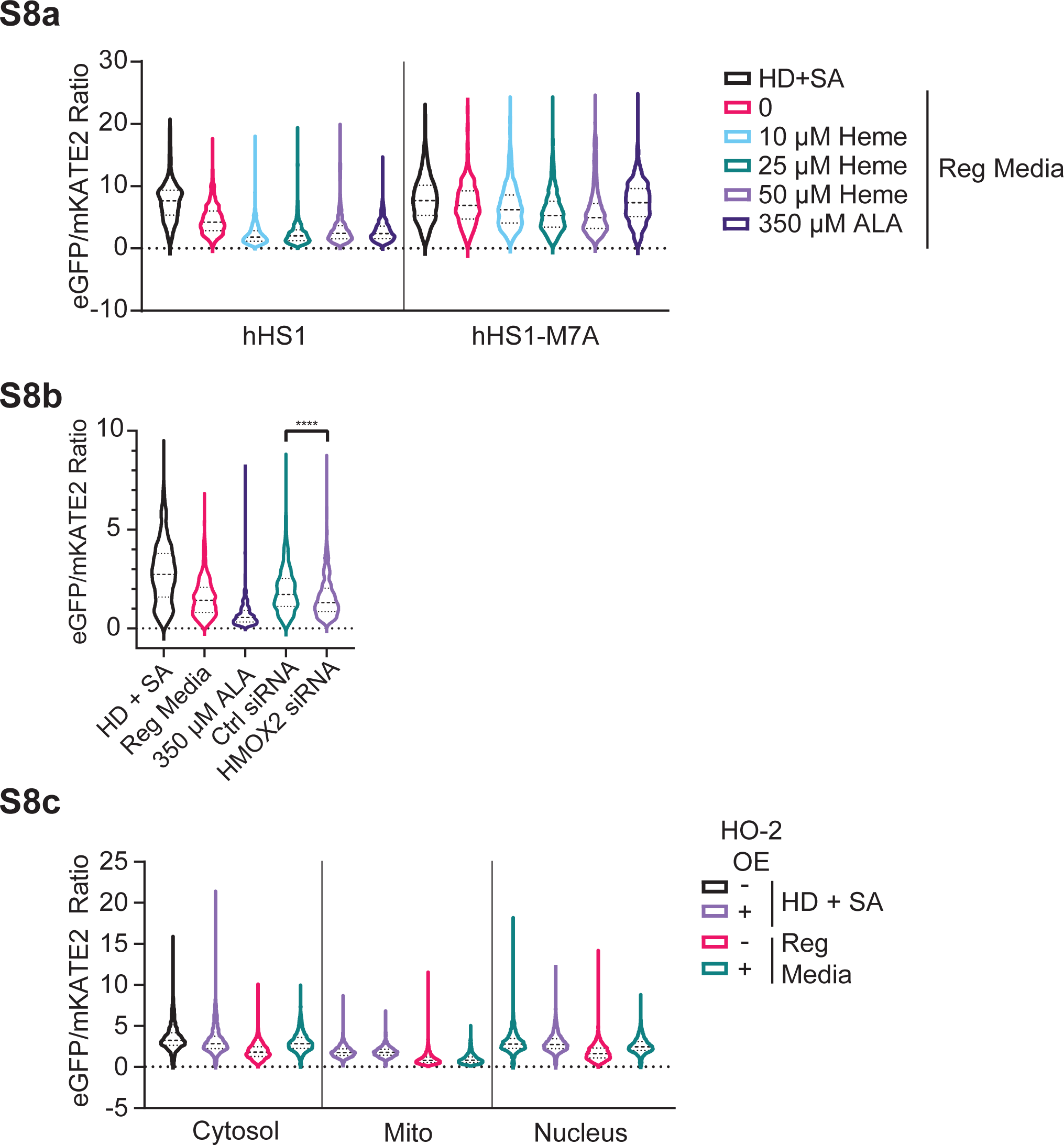

**Figure S9.**
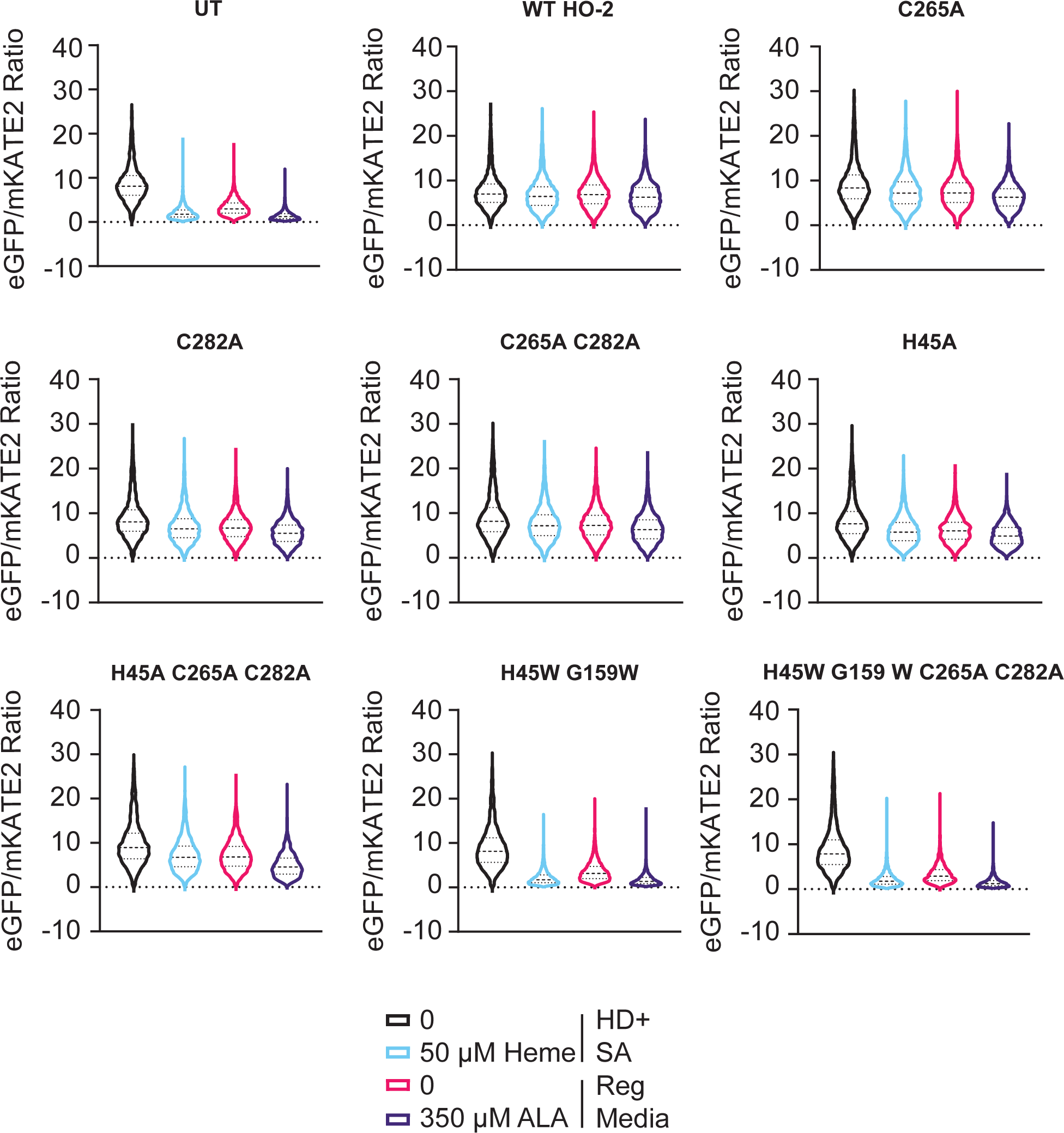

