## Supplementary Information and Figures for "Heme oxygenase-2 (HO-2) binds and buffers labile heme, which is largely oxidized, in human embryonic kidney cells"

^Co-equal authorship

**Corresponding Author (*):**

Amit R. Reddi

Georgia Institute of Technology

School of Chemistry and Biochemistry

901 Atlantic Drive, MC0400

Atlanta, GA 30332, USA

**Contents**

1. Heme Sensor Plasmid Maps and Sequences for pEF-HS1 Constructs.
2. Competition Binding Model Derivation
3. Supporting Figures (S1 – S9)
4. Supporting References

**1. Heme Sensor Plasmid Maps and Sequences for pEF5-HS1 Constructs**

**hHS1**

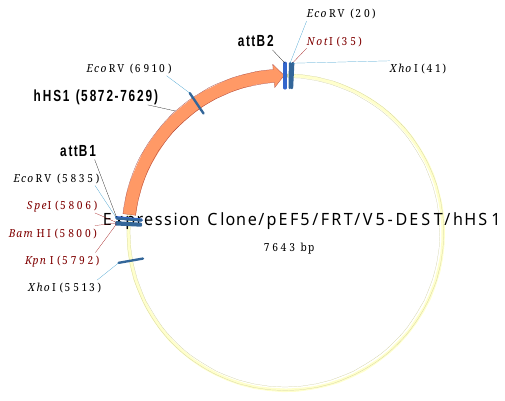

**hHS1**

ATGCACATGGTCAGCGAGCTGATCAAGGAAAACATGCACATGAAACTGTACATGGAGGGGACTGTGAACAATCACCATTTCAAATGCACCTCCGAGGGCGAAGGGAAGCCCTACGAGGGCACACAGACTATGAGGATCAAGGCAGTGGAGGGAGGACCACTGCCATTCGCCTTTGACATTCTGGCTACCTCATTCATGTACGGCAGCAAAACCTTCATCAATCACACTCAGGGGATTCCCGACTTCTTTAAGCAGTCTTTCCCTGAAGGCTTTACTTGGGAGCGAGTGACCACATACGAGGATGGAGGCGTCCTGACCGCCACACAGGACACAAGTCTGCAGGATGGCTGTCTGATCTATAACGTGAAGATTCGCGGGGTCAACTTTCCCAGTAATGGACCTGTGATGCAGAAGAAAACCCTGGGATGGGAGGCTTCAACTGAAACCCTGTACCCAGCAGACGGAGGACTGGAGGGACGAGCAGATATGGCTCTGAAACTGGTGGGCGGGGGACACCTGATCTGCAACCTGAAGACTACCTATCGGTCCAAGAAACCTGCTAAGAATCTGAAAATGCCAGGCGTGTACTATGTGGACCGGAGACTGGAGAGAATTAAGGAAGCAGATAAAGAGACCTACGTGGAGCAGCACGAAGTGGCTGTCGCACGATATTGTGACCTGCCTTCTAAACTGGGCCATCGGGGCGGGTCTATGGTGAGTAAGGGCGAGGAACTGTTCACAGGGGTGGTCCCAATCCTGGTGGAACTGGACGGCGATGTCAATGGGCACAAGTTCAGCGTGTCCGGAGAGGGAGAAGGGGACGCAACCTTTGGAGGCAGCGCCGACCTGGAAGATAATATGGAGACACTGAACGATAATCTGAAAGTGATCGAGAAAGCCGACAACGCCGCTCAGGTCAAGGATGCTCTGACTAAAATGAGGGCAGCCGCTCTGGATGCACAGAAAGCCACCCCCCCTAAGCTGGAAGACAAATCACCTGATAGCCCAGAGATGAAGGACTTCCGCCACGGATTTGATATCCTGGTCGGCCAGATTGACGATGCTCTGAAGCTGGCAAATGAAGGCAAGGTGAAAGAGGCACAGGCAGCCGCTGAGCAGCTGAAAACAACTAGGAACGCCTACCATCAGAAGTATCGCGGGGGAAAGCTGACACTGAAATTCATCTGCACCACAGGCAAGCTGCCCGTGCCCTGGCCAACTCTGGTCACTACCCTGGGATACGGCGTGCAGTGTTTTTCCCGCTATCCAGACCACATGAAGCAGCATGATTTCTTTAAATCTGCCATGCCCGAAGGCTACGTGCAGGAGAGAACCATCTTCTTTAAGGACGATGGAAACTATAAAACAAGGGCTGAAGTGAAGTTCGAGGGAGACACTCTGGTCAACCGCATCGAACTGAAGGGCATTGACTTTAAAGAGGATGGAAATATTCTGGGCCACAAGCTGGAATACAACTATAATAGCCATAACGTGTACATCATGGCCGATAAGCAGAAAAACGGCATTAAGGTCAATTTCAAAATCCGGCACAATATTGAGGACGGGAGCGTGCAGCTGGCCGATCATTACCAGCAGAACACCCCAATCGGGGACGGACCAGTGCTGCTGCCCGATAATCACTATCTGTCCACACAGTCTGCCCTGAGTAAGGACCCTAACGAAAAAAGAGATCACATGGTGCTGCTGGAGTTTGTCACCGCAGCCGGGATTACACTGGGAATGGACGAGCTGTACAAGTGA

**NLS-hHS1**

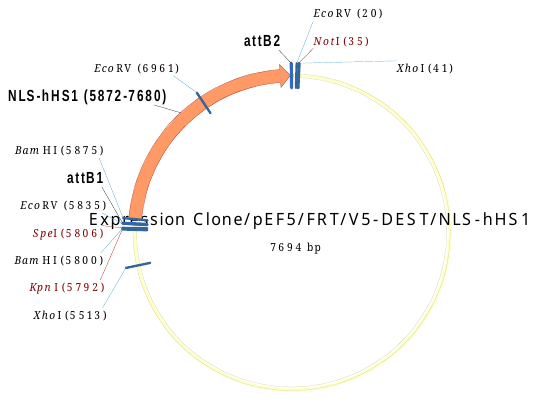

**NLS-hHS1**

ATGGATCCAAAAAAGAAGAGAAAGGTAGATCCAAAAAAGAAGAGAAAGGTAATGCACATGGTCAGCGAGCTGATCAAGGAAAACATGCACATGAAACTGTACATGGAGGGGACTGTGAACAATCACCATTTCAAATGCACCTCCGAGGGCGAAGGGAAGCCCTACGAGGGCACACAGACTATGAGGATCAAGGCAGTGGAGGGAGGACCACTGCCATTCGCCTTTGACATTCTGGCTACCTCATTCATGTACGGCAGCAAAACCTTCATCAATCACACTCAGGGGATTCCCGACTTCTTTAAGCAGTCTTTCCCTGAAGGCTTTACTTGGGAGCGAGTGACCACATACGAGGATGGAGGCGTCCTGACCGCCACACAGGACACAAGTCTGCAGGATGGCTGTCTGATCTATAACGTGAAGATTCGCGGGGTCAACTTTCCCAGTAATGGACCTGTGATGCAGAAGAAAACCCTGGGATGGGAGGCTTCAACTGAAACCCTGTACCCAGCAGACGGAGGACTGGAGGGACGAGCAGATATGGCTCTGAAACTGGTGGGCGGGGGACACCTGATCTGCAACCTGAAGACTACCTATCGGTCCAAGAAACCTGCTAAGAATCTGAAAATGCCAGGCGTGTACTATGTGGACCGGAGACTGGAGAGAATTAAGGAAGCAGATAAAGAGACCTACGTGGAGCAGCACGAAGTGGCTGTCGCACGATATTGTGACCTGCCTTCTAAACTGGGCCATCGGGGCGGGTCTATGGTGAGTAAGGGCGAGGAACTGTTCACAGGGGTGGTCCCAATCCTGGTGGAACTGGACGGCGATGTCAATGGGCACAAGTTCAGCGTGTCCGGAGAGGGAGAAGGGGACGCAACCTTTGGAGGCAGCGCCGACCTGGAAGATAATATGGAGACACTGAACGATAATCTGAAAGTGATCGAGAAAGCCGACAACGCCGCTCAGGTCAAGGATGCTCTGACTAAAATGAGGGCAGCCGCTCTGGATGCACAGAAAGCCACCCCCCCTAAGCTGGAAGACAAATCACCTGATAGCCCAGAGATGAAGGACTTCCGCCACGGATTTGATATCCTGGTCGGCCAGATTGACGATGCTCTGAAGCTGGCAAATGAAGGCAAGGTGAAAGAGGCACAGGCAGCCGCTGAGCAGCTGAAAACAACTAGGAACGCCTACCATCAGAAGTATCGCGGGGGAAAGCTGACACTGAAATTCATCTGCACCACAGGCAAGCTGCCCGTGCCCTGGCCAACTCTGGTCACTACCCTGGGATACGGCGTGCAGTGTTTTTCCCGCTATCCAGACCACATGAAGCAGCATGATTTCTTTAAATCTGCCATGCCCGAAGGCTACGTGCAGGAGAGAACCATCTTCTTTAAGGACGATGGAAACTATAAAACAAGGGCTGAAGTGAAGTTCGAGGGAGACACTCTGGTCAACCGCATCGAACTGAAGGGCATTGACTTTAAAGAGGATGGAAATATTCTGGGCCACAAGCTGGAATACAACTATAATAGCCATAACGTGTACATCATGGCCGATAAGCAGAAAAACGGCATTAAGGTCAATTTCAAAATCCGGCACAATATTGAGGACGGGAGCGTGCAGCTGGCCGATCATTACCAGCAGAACACCCCAATCGGGGACGGACCAGTGCTGCTGCCCGATAATCACTATCTGTCCACACAGTCTGCCCTGAGTAAGGACCCTAACGAAAAAAGAGATCACATGGTGCTGCTGGAGTTTGTCACCGCAGCCGGGATTACACTGGGAATGGACGAGCTGTACAAGTGA

**Mito-hHS1**

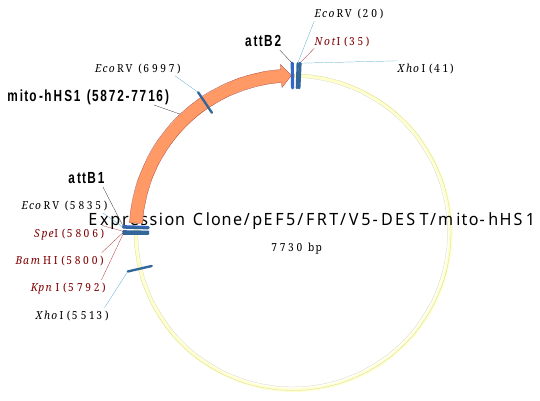

**Mito-hHS1**

ATGTCCGTCCTGACGCCGCTGCTGCTGCGGGGCTTGACAGGCTCGGCCCGGCGGCTCCCAGTGCCGCGCGCCAAGATCCATTCGTTGATGCACATGGTCAGCGAGCTGATCAAGGAAAACATGCACATGAAACTGTACATGGAGGGGACTGTGAACAATCACCATTTCAAATGCACCTCCGAGGGCGAAGGGAAGCCCTACGAGGGCACACAGACTATGAGGATCAAGGCAGTGGAGGGAGGACCACTGCCATTCGCCTTTGACATTCTGGCTACCTCATTCATGTACGGCAGCAAAACCTTCATCAATCACACTCAGGGGATTCCCGACTTCTTTAAGCAGTCTTTCCCTGAAGGCTTTACTTGGGAGCGAGTGACCACATACGAGGATGGAGGCGTCCTGACCGCCACACAGGACACAAGTCTGCAGGATGGCTGTCTGATCTATAACGTGAAGATTCGCGGGGTCAACTTTCCCAGTAATGGACCTGTGATGCAGAAGAAAACCCTGGGATGGGAGGCTTCAACTGAAACCCTGTACCCAGCAGACGGAGGACTGGAGGGACGAGCAGATATGGCTCTGAAACTGGTGGGCGGGGGACACCTGATCTGCAACCTGAAGACTACCTATCGGTCCAAGAAACCTGCTAAGAATCTGAAAATGCCAGGCGTGTACTATGTGGACCGGAGACTGGAGAGAATTAAGGAAGCAGATAAAGAGACCTACGTGGAGCAGCACGAAGTGGCTGTCGCACGATATTGTGACCTGCCTTCTAAACTGGGCCATCGGGGCGGGTCTATGGTGAGTAAGGGCGAGGAACTGTTCACAGGGGTGGTCCCAATCCTGGTGGAACTGGACGGCGATGTCAATGGGCACAAGTTCAGCGTGTCCGGAGAGGGAGAAGGGGACGCAACCTTTGGAGGCAGCGCCGACCTGGAAGATAATATGGAGACACTGAACGATAATCTGAAAGTGATCGAGAAAGCCGACAACGCCGCTCAGGTCAAGGATGCTCTGACTAAAATGAGGGCAGCCGCTCTGGATGCACAGAAAGCCACCCCCCCTAAGCTGGAAGACAAATCACCTGATAGCCCAGAGATGAAGGACTTCCGCCACGGATTTGATATCCTGGTCGGCCAGATTGACGATGCTCTGAAGCTGGCAAATGAAGGCAAGGTGAAAGAGGCACAGGCAGCCGCTGAGCAGCTGAAAACAACTAGGAACGCCTACCATCAGAAGTATCGCGGGGGAAAGCTGACACTGAAATTCATCTGCACCACAGGCAAGCTGCCCGTGCCCTGGCCAACTCTGGTCACTACCCTGGGATACGGCGTGCAGTGTTTTTCCCGCTATCCAGACCACATGAAGCAGCATGATTTCTTTAAATCTGCCATGCCCGAAGGCTACGTGCAGGAGAGAACCATCTTCTTTAAGGACGATGGAAACTATAAAACAAGGGCTGAAGTGAAGTTCGAGGGAGACACTCTGGTCAACCGCATCGAACTGAAGGGCATTGACTTTAAAGAGGATGGAAATATTCTGGGCCACAAGCTGGAATACAACTATAATAGCCATAACGTGTACATCATGGCCGATAAGCAGAAAAACGGCATTAAGGTCAATTTCAAAATCCGGCACAATATTGAGGACGGGAGCGTGCAGCTGGCCGATCATTACCAGCAGAACACCCCAATCGGGGACGGACCAGTGCTGCTGCCCGATAATCACTATCTGTCCACACAGTCTGCCCTGAGTAAGGACCCTAACGAAAAAAGAGATCACATGGTGCTGCTGGAGTTTGTCACCGCAGCCGGGATTACACTGGGAATGGACGAGCTGTACAAGTGA

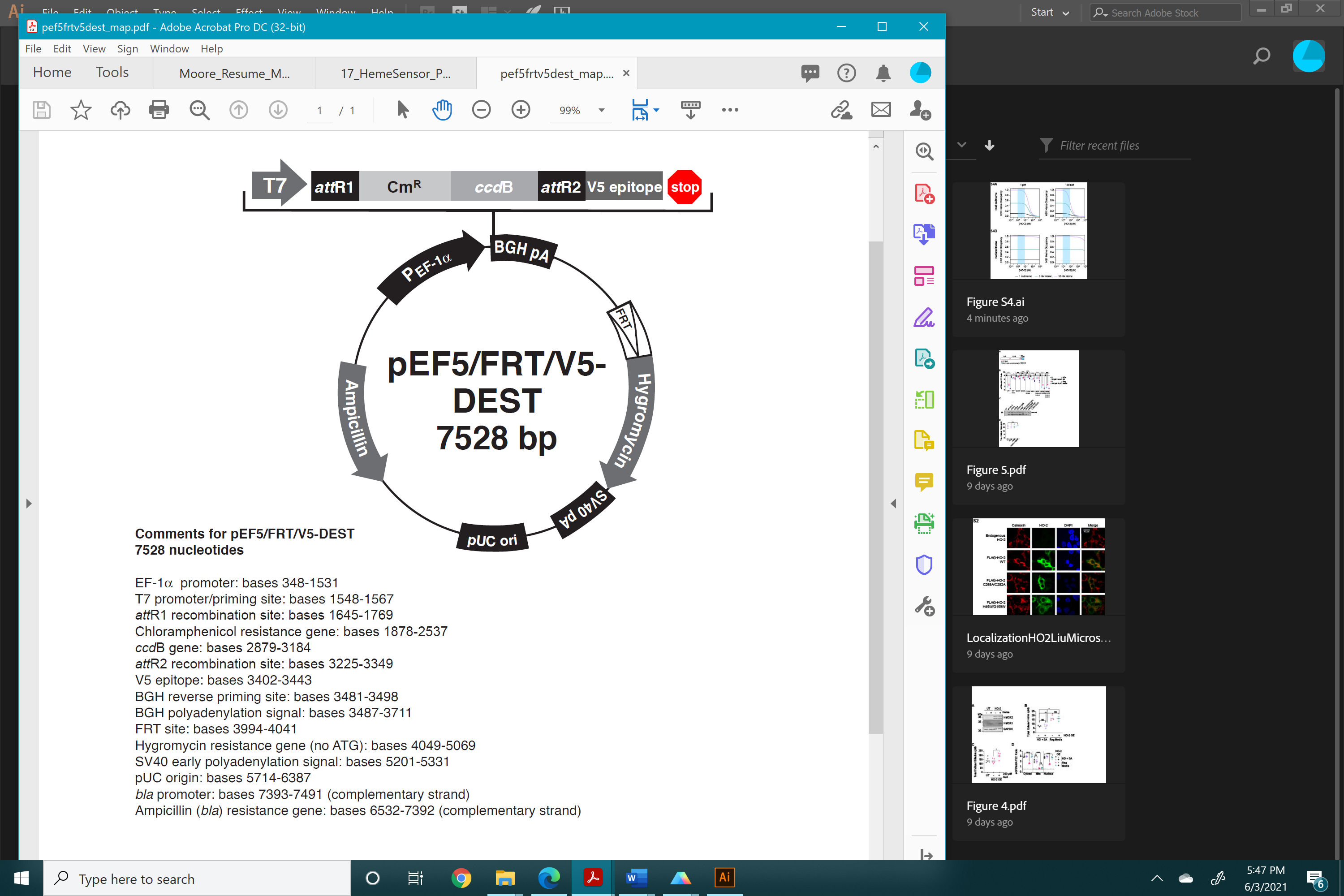

**Expression Clone pEF5FRTV5-DEST-hHS1**

cttgtacaaagtggttgatatccagcacagtggcggccgctcgagtctagagggcccgcggttcgaaggtaagcctatccctaaccctctcctcggtctcgattctacgcgtaccggttagtaatgagtttaaacccgctgatcagcctcgactgtgccttctagttgccagccatctgttgtttgcccctcccccgtgccttccttgaccctggaaggtgccactcccactgtcctttcctaataaaatgaggaaattgcatcgcattgtctgagtaggtgtcattctattctggggggtggggtggggcaggacagcaagggggaggattgggaagacaatagcaggcatgctggggatgcggtgggctctatggcttctgaggcggaaagaaccagctggggctctagggggtatccccacgcgccctgtagcggcgcattaagcgcggcgggtgtggtggttacgcgcagcgtgaccgctacacttgccagcgccctagcgcccgctcctttcgctttcttcccttcctttctcgccacgttcgccggctttccccgtcaagctctaaatcgggggtccctttagggttccgatttagtgctttacggcacctcgaccccaaaaaacttgattagggtgatggttcacgtacctagaagttcctattccgaagttcctattctctagaaagtataggaacttccttggccaaaaagcctgaactcaccgcgacgtctgtcgagaagtttctgatcgaaaagttcgacagcgtctccgacctgatgcagctctcggagggcgaagaatctcgtgctttcagcttcgatgtaggagggcgtggatatgtcctgcgggtaaatagctgcgccgatggtttctacaaagatcgttatgtttatcggcactttgcatcggccgcgctcccgattccggaagtgcttgacattggggaattcagcgagagcctgacctattgcatctcccgccgtgcacagggtgtcacgttgcaagacctgcctgaaaccgaactgcccgctgttctgcagccggtcgcggaggccatggatgcgatcgctgcggccgatcttagccagacgagcgggttcggcccattcggaccgcaaggaatcggtcaatacactacatggcgtgatttcatatgcgcgattgctgatccccatgtgtatcactggcaaactgtgatggacgacaccgtcagtgcgtccgtcgcgcaggctctcgatgagctgatgctttgggccgaggactgccccgaagtccggcacctcgtgcacgcggatttcggctccaacaatgtcctgacggacaatggccgcataacagcggtcattgactggagcgaggcgatgttcggggattcccaatacgaggtcgccaacatcttcttctggaggccgtggttggcttgtatggagcagcagacgcgctacttcgagcggaggcatccggagcttgcaggatcgccgcggctccgggcgtatatgctccgcattggtcttgaccaactctatcagagcttggttgacggcaatttcgatgatgcagcttgggcgcagggtcgatgcgacgcaatcgtccgatccggagccgggactgtcgggcgtacacaaatcgcccgcagaagcgcggccgtctggaccgatggctgtgtagaagtactcgccgatagtggaaaccgacgccccagcactcgtccgagggcaaaggaatagcacgtactacgagatttcgattccaccgccgccttctatgaaaggttgggcttcggaatcgttttccgggacgccggctggatgatcctccagcgcggggatctcatgctggagttcttcgcccaccccaacttgtttattgcagcttataatggttacaaataaagcaatagcatcacaaatttcacaaataaagcatttttttcactgcattctagttgtggtttgtccaaactcatcaatgtatcttatcatgtctgtataccgtcgacctctagctagagcttggcgtaatcatggtcatagctgtttcctgtgtgaaattgttatccgctcacaattccacacaacatacgagccggaagcataaagtgtaaagcctggggtgcctaatgagtgagctaactcacattaattgcgttgcgctcactgcccgctttccagtcgggaaacctgtcgtgccagctgcattaatgaatcggccaacgcgcggggagaggcggtttgcgtattgggcgctcttccgcttcctcgctcactgactcgctgcgctcggtcgttcggctgcggcgagcggtatcagctcactcaaaggcggtaatacggttatccacagaatcaggggataacgcaggaaagaacatgtgagcaaaaggccagcaaaaggccaggaaccgtaaaaaggccgcgttgctggcgtttttccataggctccgcccccctgacgagcatcacaaaaatcgacgctcaagtcagaggtggcgaaacccgacaggactataaagataccaggcgtttccccctggaagctccctcgtgcgctctcctgttccgaccctgccgcttaccggatacctgtccgcctttctcccttcgggaagcgtggcgctttctcatagctcacgctgtaggtatctcagttcggtgtaggtcgttcgctccaagctgggctgtgtgcacgaaccccccgttcagcccgaccgctgcgccttatccggtaactatcgtcttgagtccaacccggtaagacacgacttatcgccactggcagcagccactggtaacaggattagcagagcgaggtatgtaggcggtgctacagagttcttgaagtggtggcctaactacggctacactagaaggacagtatttggtatctgcgctctgctgaagccagttaccttcggaaaaagagttggtagctcttgatccggcaaacaaaccaccgctggtagcggtggtttttttgtttgcaagcagcagattacgcgcagaaaaaaaggatctcaagaagatcctttgatcttttctacggggtctgacgctcagtggaacgaaaactcacgttaagggattttggtcatgagattatcaaaaaggatcttcacctagatccttttaaattaaaaatgaagttttaaatcaatctaaagtatatatgagtaaacttggtctgacagttaccaatgcttaatcagtgaggcacctatctcagcgatctgtctatttcgttcatccatagttgcctgactccccgtcgtgtagataactacgatacgggagggcttaccatctggccccagtgctgcaatgataccgcgagacccacgctcaccggctccagatttatcagcaataaaccagccagccggaagggccgagcgcagaagtggtcctgcaactttatccgcctccatccagtctattaattgttgccgggaagctagagtaagtagttcgccagttaatagtttgcgcaacgttgttgccattgctacaggcatcgtggtgtcacgctcgtcgtttggtatggcttcattcagctccggttcccaacgatcaaggcgagttacatgatcccccatgttgtgcaaaaaagcggttagctccttcggtcctccgatcgttgtcagaagtaagttggccgcagtgttatcactcatggttatggcagcactgcataattctcttactgtcatgccatccgtaagatgcttttctgtgactggtgagtactcaaccaagtcattctgagaatagtgtatgcggcgaccgagttgctcttgcccggcgtcaatacgggataataccgcgccacatagcagaactttaaaagtgctcatcattggaaaacgttcttcggggcgaaaactctcaaggatcttaccgctgttgagatccagttcgatgtaacccactcgtgcacccaactgatcttcagcatcttttactttcaccagcgtttctgggtgagcaaaaacaggaaggcaaaatgccgcaaaaaagggaataagggcgacacggaaatgttgaatactcatactcttcctttttcaatattattgaagcatttatcagggttattgtctcatgagcggatacatatttgaatgtatttagaaaaataaacaaataggggttccgcgcacatttccccgaaaagtgccacctgacgtcgacggatcgggagatctcccgatcccctatggtgcactctcagtacaatctgctctgatgccgcatagttaagccagtatctgctccctgcttgtgtgttggaggtcgctgagtagtgcgcgagcaaaatttaagctacaacaaggcaaggcttgaccgacaattgcatgaagaatctgcttagggttaggcgttttgcgctgcttcgcgatgtacgggccagatatacgcgttgacattgattattgactaggcttttgcaaaaagctttgcaaagatggataaagttttaaacagagaggaatctttgcagctaatggaccttctaggtcttgaaaggagtgcctcgtgaggctccggtgcccgtcagtgggcagagcgcacatcgcccacagtccccgagaagttggggggaggggtcggcaattgaaccggtgcctagagaaggtggcgcggggtaaactgggaaagtgatgtcgtgtactggctccgcctttttcccgagggtgggggagaaccgtatataagtgcagtagtcgccgtgaacgttctttttcgcaacgggtttgccgccagaacacaggtaagtgccgtgtgtggttcccgcgggcctggcctctttacgggttatggcccttgcgtgccttgaattacttccacctggctgcagtacgtgattcttgatcccgagcttcgggttggaagtgggtgggagagttcgaggccttgcgcttaaggagccccttcgcctcgtgcttgagttgaggcctggcctgggcgctggggccgccgcgtgcgaatctggtggcaccttcgcgcctgtctcgctgctttcgataagtctctagccatttaaaatttttgatgacctgctgcgacgctttttttctggcaagatagtcttgtaaatgcgggccaagatctgcacactggtatttcggtttttggggccgcgggcggcgacggggcccgtgcgtcccagcgcacatgttcggcgaggcggggcctgcgagcgcggccaccgagaatcggacgggggtagtctcaagctggccggcctgctctggtgcctggcctcgcgccgccgtgtatcgccccgccctgggcggcaaggctggcccggtcggcaccagttgcgtgagcggaaagatggccgcttcccggccctgctgcagggagctcaaaatggaggacgcggcgctcgggagagcgggcgggtgagtcacccacacaaaggaaaagggcctttccgtcctcagccgtcgcttcatgtgactccacggagtaccgggcgccgtccaggcacctcgattagttctcgagcttttggagtacgtcgtctttaggttggggggaggggttttatgcgatggagtttccccacactgagtgggtggagactgaagttaggccagcttggcacttgatgtaattctccttggaatttgccctttttgagtttggatcttggttcattctcaagcctcagacagtggttcaaagtttttttcttccatttcaggtgtcgtgaggaattagcttggtactaatacgactcactatagggagacccaagctggctaggtaagcttggtaccgagctcggatccactagtccagtgtggtggaattctgcagatatcaacaagtttgtacaaaaaagcaggctccgccaccatgcacatggtcagcgagctgatcaaggaaaacatgcacatgaaactgtacatggaggggactgtgaacaatcaccatttcaaatgcacctccgagggcgaagggaagccctacgagggcacacagactatgaggatcaaggcagtggagggaggaccactgccattcgcctttgacattctggctacctcattcatgtacggcagcaaaaccttcatcaatcacactcaggggattcccgacttctttaagcagtctttccctgaaggctttacttgggagcgagtgaccacatacgaggatggaggcgtcctgaccgccacacaggacacaagtctgcaggatggctgtctgatctataacgtgaagattcgcggggtcaactttcccagtaatggacctgtgatgcagaagaaaaccctgggatgggaggcttcaactgaaaccctgtacccagcagacggaggactggagggacgagcagatatggctctgaaactggtgggcgggggacacctgatctgcaacctgaagactacctatcggtccaagaaacctgctaagaatctgaaaatgccaggcgtgtactatgtggaccggagactggagagaattaaggaagcagataaagagacctacgtggagcagcacgaagtggctgtcgcacgatattgtgacctgccttctaaactgggccatcggggcgggtctatggtgagtaagggcgaggaactgttcacaggggtggtcccaatcctggtggaactggacggcgatgtcaatgggcacaagttcagcgtgtccggagagggagaaggggacgcaacctttggaggcagcgccgacctggaagataatatggagacactgaacgataatctgaaagtgatcgagaaagccgacaacgccgctcaggtcaaggatgctctgactaaaatgagggcagccgctctggatgcacagaaagccaccccccctaagctggaagacaaatcacctgatagcccagagatgaaggacttccgccacggatttgatatcctggtcggccagattgacgatgctctgaagctggcaaatgaaggcaaggtgaaagaggcacaggcagccgctgagcagctgaaaacaactaggaacgcctaccatcagaagtatcgcgggggaaagctgacactgaaattcatctgcaccacaggcaagctgcccgtgccctggccaactctggtcactaccctgggatacggcgtgcagtgtttttcccgctatccagaccacatgaagcagcatgatttctttaaatctgccatgcccgaaggctacgtgcaggagagaaccatcttctttaaggacgatggaaactataaaacaagggctgaagtgaagttcgagggagacactctggtcaaccgcatcgaactgaagggcattgactttaaagaggatggaaatattctgggccacaagctggaatacaactataatagccataacgtgtacatcatggccgataagcagaaaaacggcattaaggtcaatttcaaaatccggcacaatattgaggacgggagcgtgcagctggccgatcattaccagcagaacaccccaatcggggacggaccagtgctgctgcccgataatcactatctgtccacacagtctgccctgagtaaggaccctaacgaaaaaagagatcacatggtgctgctggagtttgtcaccgcagccgggattacactgggaatggacgagctgtacaagtgagacccagcttt

**Expression Clone pEF5FRTV5-DESTNLS-hHS1**

cttgtacaaagtggttgatatccagcacagtggcggccgctcgagtctagagggcccgcggttcgaaggtaagcctatccctaaccctctcctcggtctcgattctacgcgtaccggttagtaatgagtttaaacccgctgatcagcctcgactgtgccttctagttgccagccatctgttgtttgcccctcccccgtgccttccttgaccctggaaggtgccactcccactgtcctttcctaataaaatgaggaaattgcatcgcattgtctgagtaggtgtcattctattctggggggtggggtggggcaggacagcaagggggaggattgggaagacaatagcaggcatgctggggatgcggtgggctctatggcttctgaggcggaaagaaccagctggggctctagggggtatccccacgcgccctgtagcggcgcattaagcgcggcgggtgtggtggttacgcgcagcgtgaccgctacacttgccagcgccctagcgcccgctcctttcgctttcttcccttcctttctcgccacgttcgccggctttccccgtcaagctctaaatcgggggtccctttagggttccgatttagtgctttacggcacctcgaccccaaaaaacttgattagggtgatggttcacgtacctagaagttcctattccgaagttcctattctctagaaagtataggaacttccttggccaaaaagcctgaactcaccgcgacgtctgtcgagaagtttctgatcgaaaagttcgacagcgtctccgacctgatgcagctctcggagggcgaagaatctcgtgctttcagcttcgatgtaggagggcgtggatatgtcctgcgggtaaatagctgcgccgatggtttctacaaagatcgttatgtttatcggcactttgcatcggccgcgctcccgattccggaagtgcttgacattggggaattcagcgagagcctgacctattgcatctcccgccgtgcacagggtgtcacgttgcaagacctgcctgaaaccgaactgcccgctgttctgcagccggtcgcggaggccatggatgcgatcgctgcggccgatcttagccagacgagcgggttcggcccattcggaccgcaaggaatcggtcaatacactacatggcgtgatttcatatgcgcgattgctgatccccatgtgtatcactggcaaactgtgatggacgacaccgtcagtgcgtccgtcgcgcaggctctcgatgagctgatgctttgggccgaggactgccccgaagtccggcacctcgtgcacgcggatttcggctccaacaatgtcctgacggacaatggccgcataacagcggtcattgactggagcgaggcgatgttcggggattcccaatacgaggtcgccaacatcttcttctggaggccgtggttggcttgtatggagcagcagacgcgctacttcgagcggaggcatccggagcttgcaggatcgccgcggctccgggcgtatatgctccgcattggtcttgaccaactctatcagagcttggttgacggcaatttcgatgatgcagcttgggcgcagggtcgatgcgacgcaatcgtccgatccggagccgggactgtcgggcgtacacaaatcgcccgcagaagcgcggccgtctggaccgatggctgtgtagaagtactcgccgatagtggaaaccgacgccccagcactcgtccgagggcaaaggaatagcacgtactacgagatttcgattccaccgccgccttctatgaaaggttgggcttcggaatcgttttccgggacgccggctggatgatcctccagcgcggggatctcatgctggagttcttcgcccaccccaacttgtttattgcagcttataatggttacaaataaagcaatagcatcacaaatttcacaaataaagcatttttttcactgcattctagttgtggtttgtccaaactcatcaatgtatcttatcatgtctgtataccgtcgacctctagctagagcttggcgtaatcatggtcatagctgtttcctgtgtgaaattgttatccgctcacaattccacacaacatacgagccggaagcataaagtgtaaagcctggggtgcctaatgagtgagctaactcacattaattgcgttgcgctcactgcccgctttccagtcgggaaacctgtcgtgccagctgcattaatgaatcggccaacgcgcggggagaggcggtttgcgtattgggcgctcttccgcttcctcgctcactgactcgctgcgctcggtcgttcggctgcggcgagcggtatcagctcactcaaaggcggtaatacggttatccacagaatcaggggataacgcaggaaagaacatgtgagcaaaaggccagcaaaaggccaggaaccgtaaaaaggccgcgttgctggcgtttttccataggctccgcccccctgacgagcatcacaaaaatcgacgctcaagtcagaggtggcgaaacccgacaggactataaagataccaggcgtttccccctggaagctccctcgtgcgctctcctgttccgaccctgccgcttaccggatacctgtccgcctttctcccttcgggaagcgtggcgctttctcatagctcacgctgtaggtatctcagttcggtgtaggtcgttcgctccaagctgggctgtgtgcacgaaccccccgttcagcccgaccgctgcgccttatccggtaactatcgtcttgagtccaacccggtaagacacgacttatcgccactggcagcagccactggtaacaggattagcagagcgaggtatgtaggcggtgctacagagttcttgaagtggtggcctaactacggctacactagaaggacagtatttggtatctgcgctctgctgaagccagttaccttcggaaaaagagttggtagctcttgatccggcaaacaaaccaccgctggtagcggtggtttttttgtttgcaagcagcagattacgcgcagaaaaaaaggatctcaagaagatcctttgatcttttctacggggtctgacgctcagtggaacgaaaactcacgttaagggattttggtcatgagattatcaaaaaggatcttcacctagatccttttaaattaaaaatgaagttttaaatcaatctaaagtatatatgagtaaacttggtctgacagttaccaatgcttaatcagtgaggcacctatctcagcgatctgtctatttcgttcatccatagttgcctgactccccgtcgtgtagataactacgatacgggagggcttaccatctggccccagtgctgcaatgataccgcgagacccacgctcaccggctccagatttatcagcaataaaccagccagccggaagggccgagcgcagaagtggtcctgcaactttatccgcctccatccagtctattaattgttgccgggaagctagagtaagtagttcgccagttaatagtttgcgcaacgttgttgccattgctacaggcatcgtggtgtcacgctcgtcgtttggtatggcttcattcagctccggttcccaacgatcaaggcgagttacatgatcccccatgttgtgcaaaaaagcggttagctccttcggtcctccgatcgttgtcagaagtaagttggccgcagtgttatcactcatggttatggcagcactgcataattctcttactgtcatgccatccgtaagatgcttttctgtgactggtgagtactcaaccaagtcattctgagaatagtgtatgcggcgaccgagttgctcttgcccggcgtcaatacgggataataccgcgccacatagcagaactttaaaagtgctcatcattggaaaacgttcttcggggcgaaaactctcaaggatcttaccgctgttgagatccagttcgatgtaacccactcgtgcacccaactgatcttcagcatcttttactttcaccagcgtttctgggtgagcaaaaacaggaaggcaaaatgccgcaaaaaagggaataagggcgacacggaaatgttgaatactcatactcttcctttttcaatattattgaagcatttatcagggttattgtctcatgagcggatacatatttgaatgtatttagaaaaataaacaaataggggttccgcgcacatttccccgaaaagtgccacctgacgtcgacggatcgggagatctcccgatcccctatggtgcactctcagtacaatctgctctgatgccgcatagttaagccagtatctgctccctgcttgtgtgttggaggtcgctgagtagtgcgcgagcaaaatttaagctacaacaaggcaaggcttgaccgacaattgcatgaagaatctgcttagggttaggcgttttgcgctgcttcgcgatgtacgggccagatatacgcgttgacattgattattgactaggcttttgcaaaaagctttgcaaagatggataaagttttaaacagagaggaatctttgcagctaatggaccttctaggtcttgaaaggagtgcctcgtgaggctccggtgcccgtcagtgggcagagcgcacatcgcccacagtccccgagaagttggggggaggggtcggcaattgaaccggtgcctagagaaggtggcgcggggtaaactgggaaagtgatgtcgtgtactggctccgcctttttcccgagggtgggggagaaccgtatataagtgcagtagtcgccgtgaacgttctttttcgcaacgggtttgccgccagaacacaggtaagtgccgtgtgtggttcccgcgggcctggcctctttacgggttatggcccttgcgtgccttgaattacttccacctggctgcagtacgtgattcttgatcccgagcttcgggttggaagtgggtgggagagttcgaggccttgcgcttaaggagccccttcgcctcgtgcttgagttgaggcctggcctgggcgctggggccgccgcgtgcgaatctggtggcaccttcgcgcctgtctcgctgctttcgataagtctctagccatttaaaatttttgatgacctgctgcgacgctttttttctggcaagatagtcttgtaaatgcgggccaagatctgcacactggtatttcggtttttggggccgcgggcggcgacggggcccgtgcgtcccagcgcacatgttcggcgaggcggggcctgcgagcgcggccaccgagaatcggacgggggtagtctcaagctggccggcctgctctggtgcctggcctcgcgccgccgtgtatcgccccgccctgggcggcaaggctggcccggtcggcaccagttgcgtgagcggaaagatggccgcttcccggccctgctgcagggagctcaaaatggaggacgcggcgctcgggagagcgggcgggtgagtcacccacacaaaggaaaagggcctttccgtcctcagccgtcgcttcatgtgactccacggagtaccgggcgccgtccaggcacctcgattagttctcgagcttttggagtacgtcgtctttaggttggggggaggggttttatgcgatggagtttccccacactgagtgggtggagactgaagttaggccagcttggcacttgatgtaattctccttggaatttgccctttttgagtttggatcttggttcattctcaagcctcagacagtggttcaaagtttttttcttccatttcaggtgtcgtgaggaattagcttggtactaatacgactcactatagggagacccaagctggctaggtaagcttggtaccgagctcggatccactagtccagtgtggtggaattctgcagatatcaacaagtttgtacaaaaaagcaggctccgccaccatggatccaaaaaagaagagaaaggtagatccaaaaaagaagagaaaggtaatgcacatggtcagcgagctgatcaaggaaaacatgcacatgaaactgtacatggaggggactgtgaacaatcaccatttcaaatgcacctccgagggcgaagggaagccctacgagggcacacagactatgaggatcaaggcagtggagggaggaccactgccattcgcctttgacattctggctacctcattcatgtacggcagcaaaaccttcatcaatcacactcaggggattcccgacttctttaagcagtctttccctgaaggctttacttgggagcgagtgaccacatacgaggatggaggcgtcctgaccgccacacaggacacaagtctgcaggatggctgtctgatctataacgtgaagattcgcggggtcaactttcccagtaatggacctgtgatgcagaagaaaaccctgggatgggaggcttcaactgaaaccctgtacccagcagacggaggactggagggacgagcagatatggctctgaaactggtgggcgggggacacctgatctgcaacctgaagactacctatcggtccaagaaacctgctaagaatctgaaaatgccaggcgtgtactatgtggaccggagactggagagaattaaggaagcagataaagagacctacgtggagcagcacgaagtggctgtcgcacgatattgtgacctgccttctaaactgggccatcggggcgggtctatggtgagtaagggcgaggaactgttcacaggggtggtcccaatcctggtggaactggacggcgatgtcaatgggcacaagttcagcgtgtccggagagggagaaggggacgcaacctttggaggcagcgccgacctggaagataatatggagacactgaacgataatctgaaagtgatcgagaaagccgacaacgccgctcaggtcaaggatgctctgactaaaatgagggcagccgctctggatgcacagaaagccaccccccctaagctggaagacaaatcacctgatagcccagagatgaaggacttccgccacggatttgatatcctggtcggccagattgacgatgctctgaagctggcaaatgaaggcaaggtgaaagaggcacaggcagccgctgagcagctgaaaacaactaggaacgcctaccatcagaagtatcgcgggggaaagctgacactgaaattcatctgcaccacaggcaagctgcccgtgccctggccaactctggtcactaccctgggatacggcgtgcagtgtttttcccgctatccagaccacatgaagcagcatgatttctttaaatctgccatgcccgaaggctacgtgcaggagagaaccatcttctttaaggacgatggaaactataaaacaagggctgaagtgaagttcgagggagacactctggtcaaccgcatcgaactgaagggcattgactttaaagaggatggaaatattctgggccacaagctggaatacaactataatagccataacgtgtacatcatggccgataagcagaaaaacggcattaaggtcaatttcaaaatccggcacaatattgaggacgggagcgtgcagctggccgatcattaccagcagaacaccccaatcggggacggaccagtgctgctgcccgataatcactatctgtccacacagtctgccctgagtaaggaccctaacgaaaaaagagatcacatggtgctgctggagtttgtcaccgcagccgggattacactgggaatggacgagctgtacaagtgagacccagcttt

**Expression ClonepEF5FRTV5-DESTmito-hHS1**

cttgtacaaagtggttgatatccagcacagtggcggccgctcgagtctagagggcccgcggttcgaaggtaagcctatccctaaccctctcctcggtctcgattctacgcgtaccggttagtaatgagtttaaacccgctgatcagcctcgactgtgccttctagttgccagccatctgttgtttgcccctcccccgtgccttccttgaccctggaaggtgccactcccactgtcctttcctaataaaatgaggaaattgcatcgcattgtctgagtaggtgtcattctattctggggggtggggtggggcaggacagcaagggggaggattgggaagacaatagcaggcatgctggggatgcggtgggctctatggcttctgaggcggaaagaaccagctggggctctagggggtatccccacgcgccctgtagcggcgcattaagcgcggcgggtgtggtggttacgcgcagcgtgaccgctacacttgccagcgccctagcgcccgctcctttcgctttcttcccttcctttctcgccacgttcgccggctttccccgtcaagctctaaatcgggggtccctttagggttccgatttagtgctttacggcacctcgaccccaaaaaacttgattagggtgatggttcacgtacctagaagttcctattccgaagttcctattctctagaaagtataggaacttccttggccaaaaagcctgaactcaccgcgacgtctgtcgagaagtttctgatcgaaaagttcgacagcgtctccgacctgatgcagctctcggagggcgaagaatctcgtgctttcagcttcgatgtaggagggcgtggatatgtcctgcgggtaaatagctgcgccgatggtttctacaaagatcgttatgtttatcggcactttgcatcggccgcgctcccgattccggaagtgcttgacattggggaattcagcgagagcctgacctattgcatctcccgccgtgcacagggtgtcacgttgcaagacctgcctgaaaccgaactgcccgctgttctgcagccggtcgcggaggccatggatgcgatcgctgcggccgatcttagccagacgagcgggttcggcccattcggaccgcaaggaatcggtcaatacactacatggcgtgatttcatatgcgcgattgctgatccccatgtgtatcactggcaaactgtgatggacgacaccgtcagtgcgtccgtcgcgcaggctctcgatgagctgatgctttgggccgaggactgccccgaagtccggcacctcgtgcacgcggatttcggctccaacaatgtcctgacggacaatggccgcataacagcggtcattgactggagcgaggcgatgttcggggattcccaatacgaggtcgccaacatcttcttctggaggccgtggttggcttgtatggagcagcagacgcgctacttcgagcggaggcatccggagcttgcaggatcgccgcggctccgggcgtatatgctccgcattggtcttgaccaactctatcagagcttggttgacggcaatttcgatgatgcagcttgggcgcagggtcgatgcgacgcaatcgtccgatccggagccgggactgtcgggcgtacacaaatcgcccgcagaagcgcggccgtctggaccgatggctgtgtagaagtactcgccgatagtggaaaccgacgccccagcactcgtccgagggcaaaggaatagcacgtactacgagatttcgattccaccgccgccttctatgaaaggttgggcttcggaatcgttttccgggacgccggctggatgatcctccagcgcggggatctcatgctggagttcttcgcccaccccaacttgtttattgcagcttataatggttacaaataaagcaatagcatcacaaatttcacaaataaagcatttttttcactgcattctagttgtggtttgtccaaactcatcaatgtatcttatcatgtctgtataccgtcgacctctagctagagcttggcgtaatcatggtcatagctgtttcctgtgtgaaattgttatccgctcacaattccacacaacatacgagccggaagcataaagtgtaaagcctggggtgcctaatgagtgagctaactcacattaattgcgttgcgctcactgcccgctttccagtcgggaaacctgtcgtgccagctgcattaatgaatcggccaacgcgcggggagaggcggtttgcgtattgggcgctcttccgcttcctcgctcactgactcgctgcgctcggtcgttcggctgcggcgagcggtatcagctcactcaaaggcggtaatacggttatccacagaatcaggggataacgcaggaaagaacatgtgagcaaaaggccagcaaaaggccaggaaccgtaaaaaggccgcgttgctggcgtttttccataggctccgcccccctgacgagcatcacaaaaatcgacgctcaagtcagaggtggcgaaacccgacaggactataaagataccaggcgtttccccctggaagctccctcgtgcgctctcctgttccgaccctgccgcttaccggatacctgtccgcctttctcccttcgggaagcgtggcgctttctcatagctcacgctgtaggtatctcagttcggtgtaggtcgttcgctccaagctgggctgtgtgcacgaaccccccgttcagcccgaccgctgcgccttatccggtaactatcgtcttgagtccaacccggtaagacacgacttatcgccactggcagcagccactggtaacaggattagcagagcgaggtatgtaggcggtgctacagagttcttgaagtggtggcctaactacggctacactagaaggacagtatttggtatctgcgctctgctgaagccagttaccttcggaaaaagagttggtagctcttgatccggcaaacaaaccaccgctggtagcggtggtttttttgtttgcaagcagcagattacgcgcagaaaaaaaggatctcaagaagatcctttgatcttttctacggggtctgacgctcagtggaacgaaaactcacgttaagggattttggtcatgagattatcaaaaaggatcttcacctagatccttttaaattaaaaatgaagttttaaatcaatctaaagtatatatgagtaaacttggtctgacagttaccaatgcttaatcagtgaggcacctatctcagcgatctgtctatttcgttcatccatagttgcctgactccccgtcgtgtagataactacgatacgggagggcttaccatctggccccagtgctgcaatgataccgcgagacccacgctcaccggctccagatttatcagcaataaaccagccagccggaagggccgagcgcagaagtggtcctgcaactttatccgcctccatccagtctattaattgttgccgggaagctagagtaagtagttcgccagttaatagtttgcgcaacgttgttgccattgctacaggcatcgtggtgtcacgctcgtcgtttggtatggcttcattcagctccggttcccaacgatcaaggcgagttacatgatcccccatgttgtgcaaaaaagcggttagctccttcggtcctccgatcgttgtcagaagtaagttggccgcagtgttatcactcatggttatggcagcactgcataattctcttactgtcatgccatccgtaagatgcttttctgtgactggtgagtactcaaccaagtcattctgagaatagtgtatgcggcgaccgagttgctcttgcccggcgtcaatacgggataataccgcgccacatagcagaactttaaaagtgctcatcattggaaaacgttcttcggggcgaaaactctcaaggatcttaccgctgttgagatccagttcgatgtaacccactcgtgcacccaactgatcttcagcatcttttactttcaccagcgtttctgggtgagcaaaaacaggaaggcaaaatgccgcaaaaaagggaataagggcgacacggaaatgttgaatactcatactcttcctttttcaatattattgaagcatttatcagggttattgtctcatgagcggatacatatttgaatgtatttagaaaaataaacaaataggggttccgcgcacatttccccgaaaagtgccacctgacgtcgacggatcgggagatctcccgatcccctatggtgcactctcagtacaatctgctctgatgccgcatagttaagccagtatctgctccctgcttgtgtgttggaggtcgctgagtagtgcgcgagcaaaatttaagctacaacaaggcaaggcttgaccgacaattgcatgaagaatctgcttagggttaggcgttttgcgctgcttcgcgatgtacgggccagatatacgcgttgacattgattattgactaggcttttgcaaaaagctttgcaaagatggataaagttttaaacagagaggaatctttgcagctaatggaccttctaggtcttgaaaggagtgcctcgtgaggctccggtgcccgtcagtgggcagagcgcacatcgcccacagtccccgagaagttggggggaggggtcggcaattgaaccggtgcctagagaaggtggcgcggggtaaactgggaaagtgatgtcgtgtactggctccgcctttttcccgagggtgggggagaaccgtatataagtgcagtagtcgccgtgaacgttctttttcgcaacgggtttgccgccagaacacaggtaagtgccgtgtgtggttcccgcgggcctggcctctttacgggttatggcccttgcgtgccttgaattacttccacctggctgcagtacgtgattcttgatcccgagcttcgggttggaagtgggtgggagagttcgaggccttgcgcttaaggagccccttcgcctcgtgcttgagttgaggcctggcctgggcgctggggccgccgcgtgcgaatctggtggcaccttcgcgcctgtctcgctgctttcgataagtctctagccatttaaaatttttgatgacctgctgcgacgctttttttctggcaagatagtcttgtaaatgcgggccaagatctgcacactggtatttcggtttttggggccgcgggcggcgacggggcccgtgcgtcccagcgcacatgttcggcgaggcggggcctgcgagcgcggccaccgagaatcggacgggggtagtctcaagctggccggcctgctctggtgcctggcctcgcgccgccgtgtatcgccccgccctgggcggcaaggctggcccggtcggcaccagttgcgtgagcggaaagatggccgcttcccggccctgctgcagggagctcaaaatggaggacgcggcgctcgggagagcgggcgggtgagtcacccacacaaaggaaaagggcctttccgtcctcagccgtcgcttcatgtgactccacggagtaccgggcgccgtccaggcacctcgattagttctcgagcttttggagtacgtcgtctttaggttggggggaggggttttatgcgatggagtttccccacactgagtgggtggagactgaagttaggccagcttggcacttgatgtaattctccttggaatttgccctttttgagtttggatcttggttcattctcaagcctcagacagtggttcaaagtttttttcttccatttcaggtgtcgtgaggaattagcttggtactaatacgactcactatagggagacccaagctggctaggtaagcttggtaccgagctcggatccactagtccagtgtggtggaattctgcagatatcaacaagtttgtacaaaaaagcaggctccgccaccatgtccgtcctgacgccgctgctgctgcggggcttgacaggctcggcccggcggctcccagtgccgcgcgccaagatccattcgttgatgcacatggtcagcgagctgatcaaggaaaacatgcacatgaaactgtacatggaggggactgtgaacaatcaccatttcaaatgcacctccgagggcgaagggaagccctacgagggcacacagactatgaggatcaaggcagtggagggaggaccactgccattcgcctttgacattctggctacctcattcatgtacggcagcaaaaccttcatcaatcacactcaggggattcccgacttctttaagcagtctttccctgaaggctttacttgggagcgagtgaccacatacgaggatggaggcgtcctgaccgccacacaggacacaagtctgcaggatggctgtctgatctataacgtgaagattcgcggggtcaactttcccagtaatggacctgtgatgcagaagaaaaccctgggatgggaggcttcaactgaaaccctgtacccagcagacggaggactggagggacgagcagatatggctctgaaactggtgggcgggggacacctgatctgcaacctgaagactacctatcggtccaagaaacctgctaagaatctgaaaatgccaggcgtgtactatgtggaccggagactggagagaattaaggaagcagataaagagacctacgtggagcagcacgaagtggctgtcgcacgatattgtgacctgccttctaaactgggccatcggggcgggtctatggtgagtaagggcgaggaactgttcacaggggtggtcccaatcctggtggaactggacggcgatgtcaatgggcacaagttcagcgtgtccggagagggagaaggggacgcaacctttggaggcagcgccgacctggaagataatatggagacactgaacgataatctgaaagtgatcgagaaagccgacaacgccgctcaggtcaaggatgctctgactaaaatgagggcagccgctctggatgcacagaaagccaccccccctaagctggaagacaaatcacctgatagcccagagatgaaggacttccgccacggatttgatatcctggtcggccagattgacgatgctctgaagctggcaaatgaaggcaaggtgaaagaggcacaggcagccgctgagcagctgaaaacaactaggaacgcctaccatcagaagtatcgcgggggaaagctgacactgaaattcatctgcaccacaggcaagctgcccgtgccctggccaactctggtcactaccctgggatacggcgtgcagtgtttttcccgctatccagaccacatgaagcagcatgatttctttaaatctgccatgcccgaaggctacgtgcaggagagaaccatcttctttaaggacgatggaaactataaaacaagggctgaagtgaagttcgagggagacactctggtcaaccgcatcgaactgaagggcattgactttaaagaggatggaaatattctgggccacaagctggaatacaactataatagccataacgtgtacatcatggccgataagcagaaaaacggcattaaggtcaatttcaaaatccggcacaatattgaggacgggagcgtgcagctggccgatcattaccagcagaacaccccaatcggggacggaccagtgctgctgcccgataatcactatctgtccacacagtctgccctgagtaaggaccctaacgaaaaaagagatcacatggtgctgctggagtttgtcaccgcagccgggattacactgggaatggacgagctgtacaagtgagacccagcttt

**2. Competition Binding Model Derivation**

**Competition model between HS1 and HO-2 (Figures 6 and S4):**

We modeled HS1 heme occupancy as a function of HO-2 expression for varying concentrations of heme and heme sensor. The changes in HS1 heme occupancy as a function of HO-2 expression is consistent with the bulk of labile heme being oxidized in HEK293 cells.

Using the following heme competition model and mass balance equations:

HO2-Heme + HS1 🡨🡪 HO2 + HS1-Heme

[HS1]_T_ = [HS1-Heme] + [HS1]

[Heme]_T_ = [HS1-Heme] + [HO2-Heme]

[HO2-Heme]_T_ = [HS1-Heme] + [HO2-Heme]

*K*_C_ (= *K*_D_^HS1-Heme^/*K*_D_^HO2-Heme^) = (HS1)(HO2-Heme) / (HO2)(HS1-Heme)

one can derive an equation that relates the fractional heme occupancy of HS1 to the amount of HO2, Heme, and HS1 present.

Fraction Heme Occupancy of HS1 = (-b – (b^2^-4ac)^.5^ / 2a) / HS1_T_

a = 1 - *K*_C_

b = (-Heme_T_ – HS1_T_ + *K*_C_ * Heme_T_ - *K*_C_ * HO2_T_)

c = HS1_T_ * Heme_T_

To simulate the competition, the following was assumed:

*K*_C_ was fixed as either 0.8 for ferric heme (K_D_^HS1^ = 3.0 nM and K_D_^HO-2^ = 3.6 nM) or 3.3 x 10^-6^ for ferrous heme (K_D_^HS1^ = 1 pM and K_D_^HO-2^ = 320 nM). The competition was modeled for varying levels of HS1_T_, including 10 nM (**Figure 6**), 100 nM (**Figure S4**), and 1 µM (**Figure S4**) and Heme_T_, including 0.1, 0.5, an 1.0 equivalents of heme relative to [HS1]; For reference, quantitative immunoblotting in yeast and HEK293 cells indicate that the sensor is expressed at levels spanning 1-20 nM.

**3.	Supporting Figures**

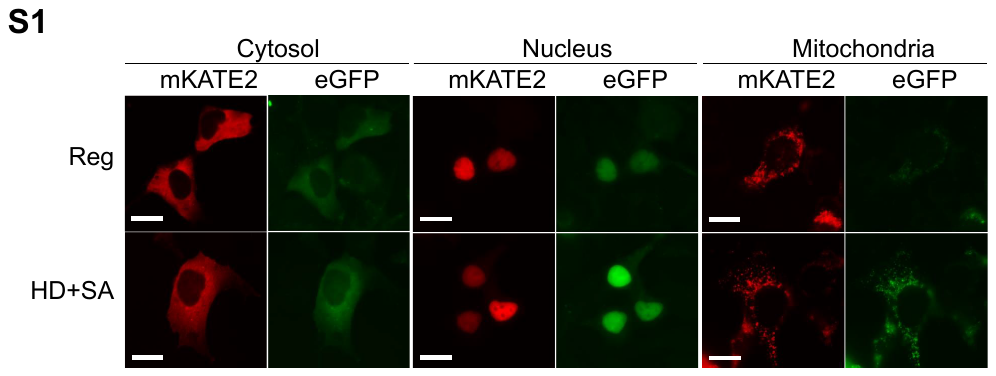

**Figure S1.** Fluorescence imaging of cytosolic, nuclear, and mitochondrial-targeted HS1. Scale bar = 10 µm.

**Figure S2.** HO2 and its variants localize to the ER. HEK293 cells cultured on coverslips were transfected with transfection reagent only (Endogenous HO2) or with corresponding expression vector for FLAG-tagged HO2 and its variants as indicated. Endogenous HO2 was stained with anti-HO2 antibody, FLAG-tagged HO2 was probed with anti-FLAG antibody. Calnexin was used as a marker for ER and DAPI was used to show the position of nuclei. “Merge” is the merged images of Calnexin channel and HO2 channel. All images are in the same scale as indicated with the 10 µm scale bar.

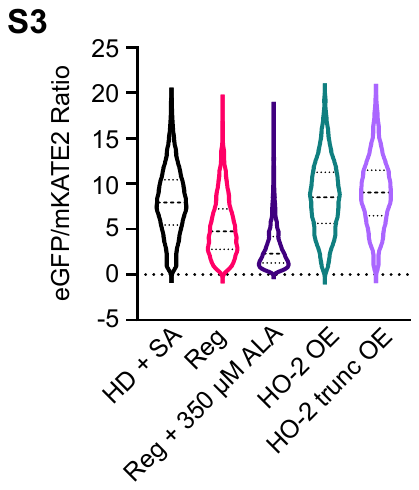

**Figure S3.** Deletion of the membrane spanning region of HO-2, 288-316 (HO-2 trunc), which tethers HO-2 to the ER membrane, does not affect the ability of HO-2 to deplete labile heme when overexpressed (OE). Cells were analyzed by flow cytometry as described in **Experimental Procedures** and cultured to be heme depleted (HD+SA), heme replete (Reg), or have excess heme (Reg + 350 µM ALA).

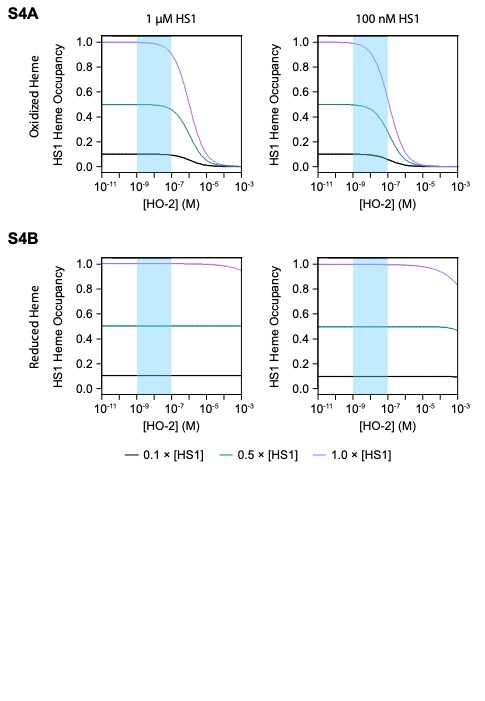

**Figure S4.** The effects of HO-2 depletion or overexpression on HS1 heme occupancy. (**a**) Simulation of HS1 heme occupancy as a function of HO-2 expression assuming the equilibrium model depicted in **Equation 3**, with [HS1] = 100 nM or 1 µM, and heme being oxidized and present at 0.1 (black), 0.5 (green), or 1.0 (pink) equivalent relative to [HS1]. (**b**) Simulation of HS1 heme occupancy as a function of HO-2 expression as in panel “a”, except assuming heme is reduced. See main text for details and **Supporting Information** for derivations of the equilibrium model. See also **Figure 6**, which depicts simulations for 10 nM [HS1].

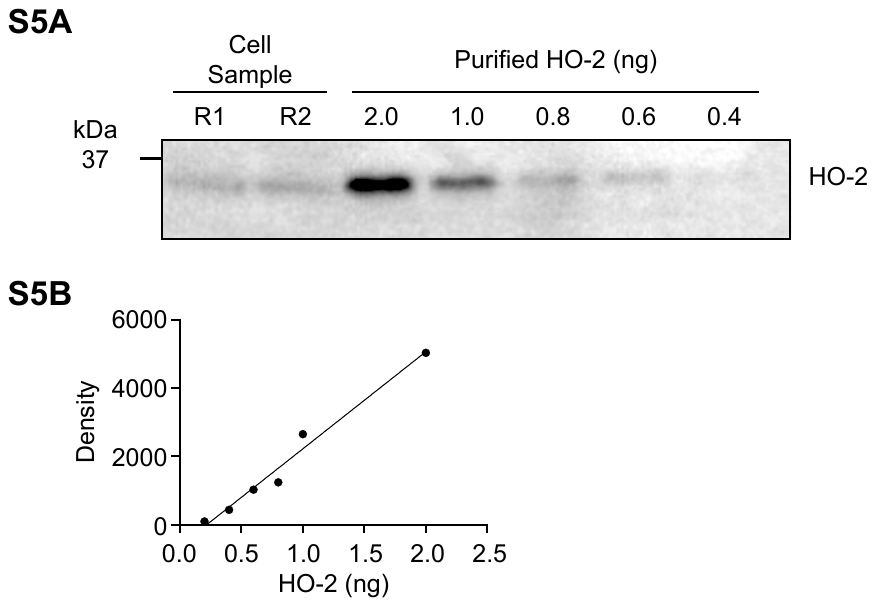

**Figure S5.** Quantitative immunoblotting to determine the concentration of endogenous HO-2 expression in HEK293 cells. Analysis reveals that HO-2 is expressed at a concentration of ~10 nM, assuming a HEK293 cell volume of 1.2 pL.

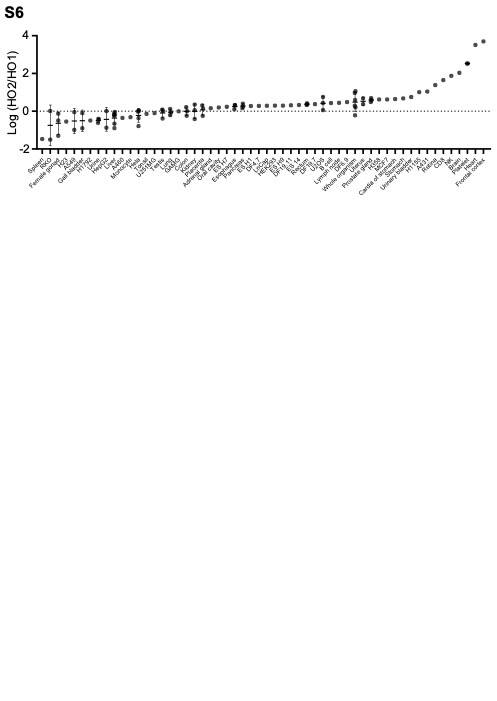

**Figure S6.** Relative HO-2 and HO-1 protein expression across various cell types and tissues as assessed by quantitative proteomics (1,2).

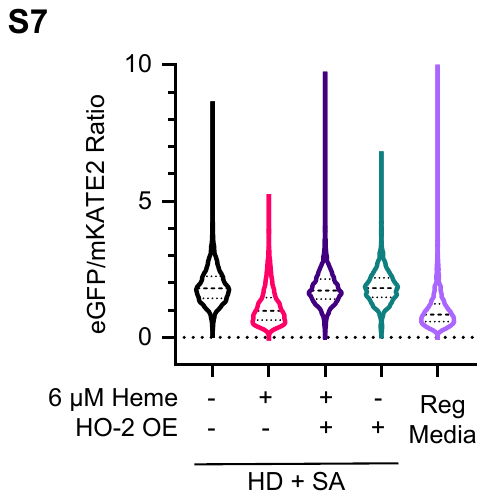

**Figure S7.** Overexpression of HO-2 is able to deplete mitochondrial labile heme if heme is supplied exogenously. Cells were analyzed by flow cytometry as described in **Experimental Procedures** and cultured to be heme depleted (HD+SA), heme replete (Reg), or supplemented with the indicated concentration of heme. This is in contrast to endogenously synthesized labile heme in the mitochondria, which is unaffected by HO-2 overexpression (**Figure 4**).

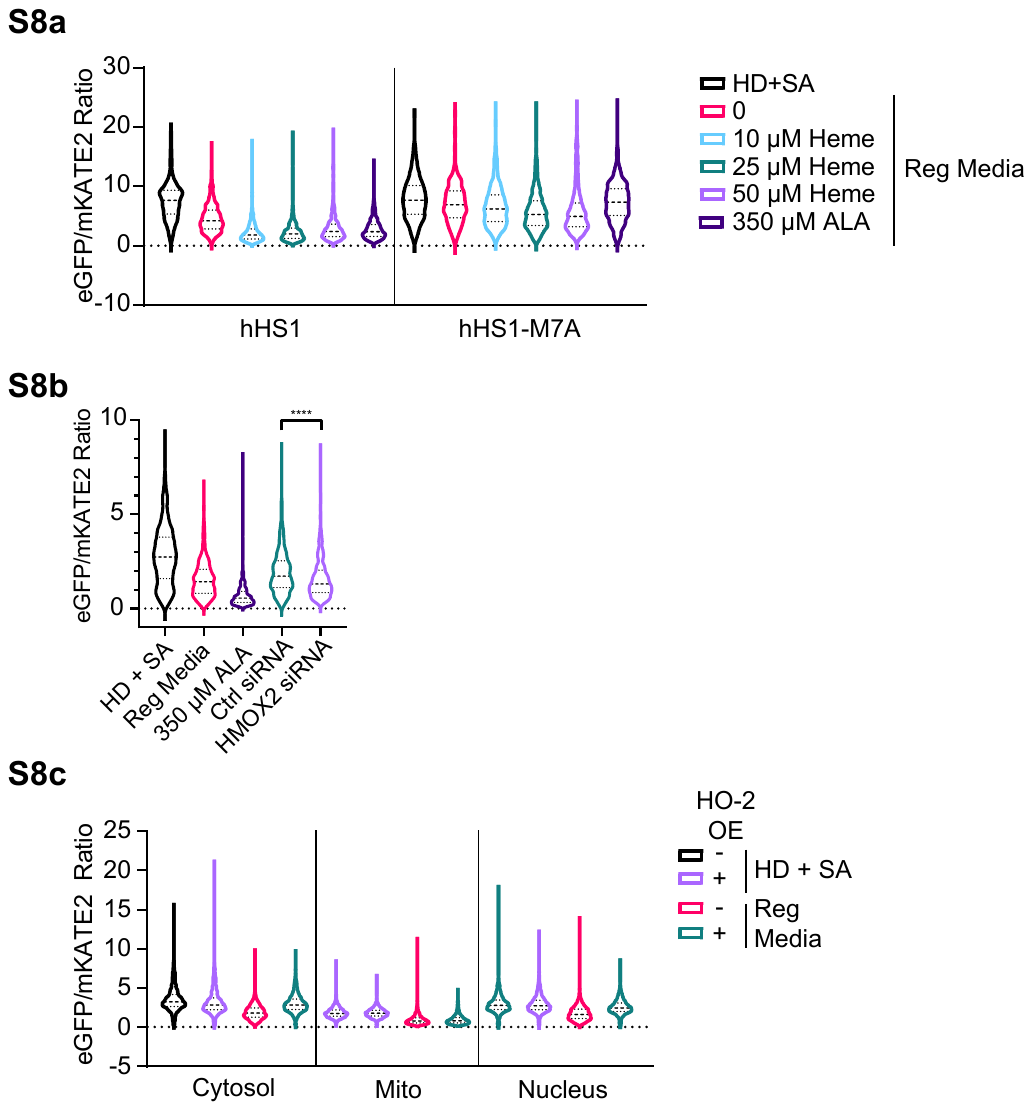

**Figure S8.** Representative flow cytometry histograms, as depicted using violin plots, of HS1 eGFP/mKATE2 fluorescence ratios in HEK293 cells expressing the indicated sensors. (**a**) Representative violin plots of heme sensor eGFP/mKATE2 fluorescence ratio distributions from single cell analysis of HEK293 cultures are shown for cells grown in HD + SA media or in regular media supplemented with the indicated concentrations of hemin chloride or 5-aminolevulinic acid (ALA) for 24 hours. See **Figure 2b**. (**b**) Representative violin plots of heme sensor eGFP/mKATE2 fluorescence ratio distributions from single cell analysis of HEK293 cultures are shown for HEK293 grown in HD + SA media, regular media, or regular media supplemented with 350 µM ALA or control or targeted siRNA against HMOX2. See **Figure 3b**. (**c**) Representative violin plots of heme sensor eGFP/mKATE2 fluorescence ratio distributions from single cell analysis of HEK293 cultures are shown for HEK293 cells expressing cytosolic, nuclear or mitochondrial (mito)-targeted HS1 in untranfected (-) or HO-2 overexpressing (OE) (+) HEK293 cells grown in HD + SA or regular media. See **Figure S8c**.

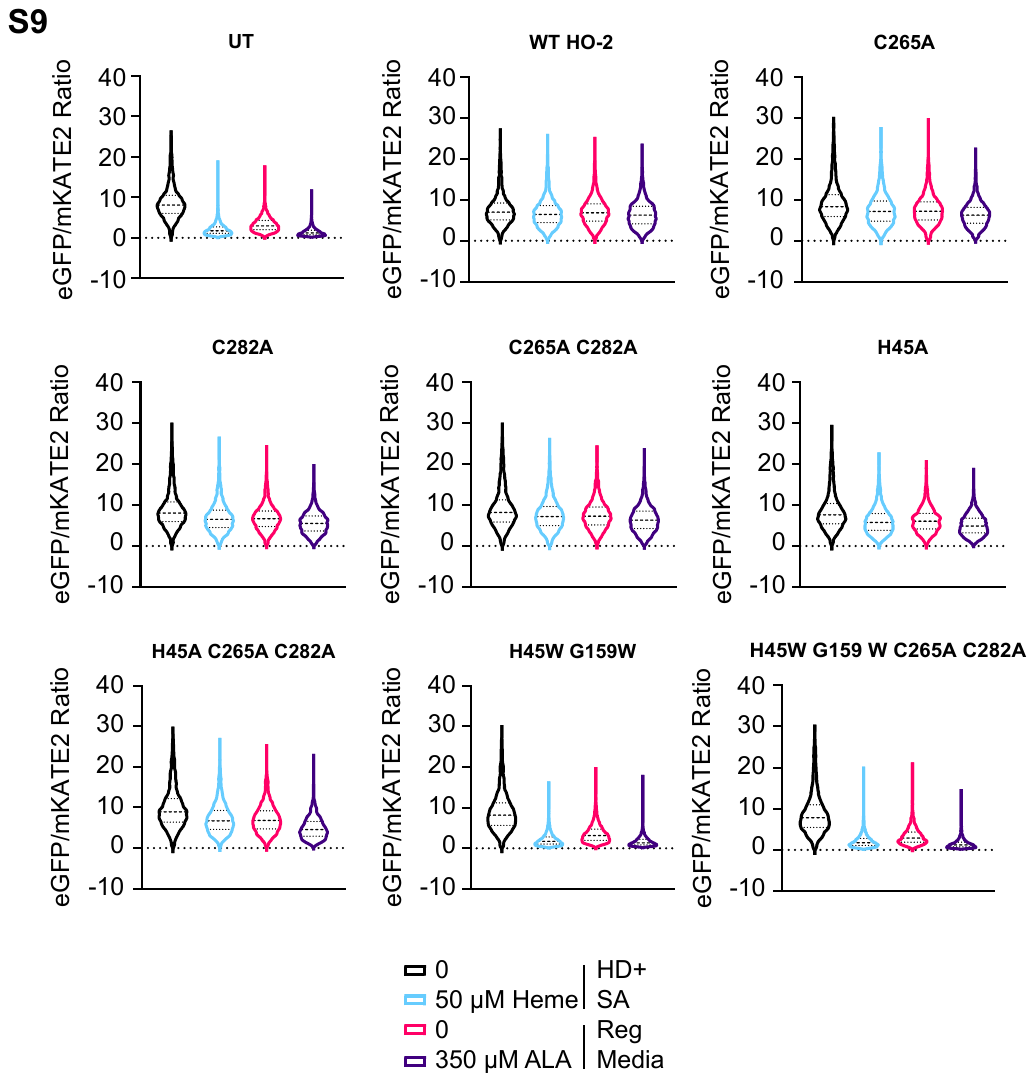

**Figure S9.** Representative flow cytometry histograms, as depicted using violin plots, of HS1 eGFP/mKATE2 fluorescence ratios in HEK293 cells expressing the indicated HO-2 variants grown in HD + SA media or in regular media supplemented with the indicated concentrations of hemin chloride or 5-aminolevulinic acid (ALA) for 24 hours. See **Figure 5b**.
